# Modelling Discrete States and Long-Term Dynamics in Functional Brain Networks

**DOI:** 10.1101/2025.09.25.678554

**Authors:** SungJun Cho, Rukuang Huang, Chetan Gohil, Oiwi Parker Jones, Mark W. Woolrich

## Abstract

Functional brain network dynamics underlie fundamental aspects of human cognition and behaviour, including memory, ageing, and a range of clinical disorders. It has been shown that ongoing brain network dynamics can be reliably inferred at fast, sub-second timescales from electrophysiological data using unsupervised machine learning. However, these methods often struggle with inherent trade-offs. For example, Hidden Markov Models (HMMs) have been used to infer categorical brain network states that provide good interpretability but do not model long-range temporal structure. Recently, deep learning approaches using recurrent neural networks (e.g., Dynamic Network Modes) have been proposed to model long-range temporal dependencies, but at the expense of interpretability. In this paper, we introduce Dynamic Network States (DyNeStE) to address this problem. This new model employs amortised Bayesian inference with recurrent neural networks to model long-range temporal structure and uses a Gumbel-Softmax distribution to enforce categorical states for greater interpretability. In both simulations and real resting-state magnetoencephalography data, DyNeStE was able to recover plausible dynamic brain network states and showed superior performance over the HMM in capturing long-range temporal dependencies in network dynamics. These dynamic networks were reproducible across independent data splits and build on established HMM-based findings. Together, these results highlight DyNeStE as an interpretable and temporally informative framework, capable of representing large-scale neural activity as discrete state transitions while capturing transient and long-range brain network dynamics.

## 1 Introduction

During rest and task, the human brain is characterised by time-varying spatiotemporal networks that activate in sub-second timescales (Baker et al., 2014; Quinn et al., 2018; Vidaurre et al., 2016). These dynamic brain networks have been observed consistently in functional neuroimaging signals. In electrophysiology, it is now well established that these networks are linked to cognition (Barnes et al., 2015; Becker et al., 2020), ageing (Gohil, Kohl, Pitt, et al., 2024; Tibon et al., 2021), and clinical disorders (Kohl et al., 2025; Rossi et al., 2024; Sitnikova et al., 2018). Moreover, the spatial and spectral profiles of the dynamic networks were found to be comparable across functional magnetic resonance imaging (fMRI), electroencephalography (EEG), and magnetoencephalography (MEG) at rest (Brookes et al., 2011; Cho et al., 2024), implying that they are reliable biological constructs that can be robustly measured across data modalities. Accurately modelling these networks, therefore, is a central challenge in the study of human cognition and behaviour.

Current state-of-the-art methods for estimating dynamic brain networks include the Hidden Markov Model (HMM) (Baker et al., 2014; Vidaurre et al., 2018) and deep learning approaches such as Dynamic Network Modes (DyNeMo) (Gohil et al., 2022). These methods use probabilistic, generative models to characterise the temporal dynamics of brain networks without imposing prior assumptions about their timings or timescales. As alternatives, traditional methods such as sliding-window analysis (Betti et al., 2013; De Pasquale et al., 2010) or Independent Component Analysis (ICA) (Brookes et al., 2011) have been widely used, but they lack probabilistic interpretations and/or rely heavily on heuristic assumptions that constrain the duration of network activations.

Existing generative models (the HMM and DyNeMo), however, have some limitations. First, HMMs assume that network states depend only on the immediately preceding state^1^, thereby struggling to capture long-range temporal dependencies in the state dynamics. On the other hand, existing neural network-based models typically represent brain network dynamics in a continuous latent space. For instance, DyNeMo assumes a time-varying linear mixture of network modes that allows multiple networks to coactivate simultaneously. This lack of a clearly defined, mutually exclusive brain state at each time point can compromise interpretability, making it difficult to identify recurrent whole-brain network activity (van Es, Higgins, et al., 2025). While the Hidden Semi-Markov Model (HSMM; Trujillo-Barreto et al., 2024) has been proposed to better capture prolonged state activations, it nevertheless remains constrained by the first-order Markovian assumption and requires parametric specification of distributions that define state durations.

To address these limitations and to provide an intermediate approach, we introduce **Dynamic Network States (DyNeStE)**, a novel generative model that learns a latent categorical description of network dynamics, i.e., a set of ‘states’ that correspond to distinct functional brain networks. The proposed model is designed based on a variational autoencoder (VAE) (Kingma & Welling, 2014) with a Gumbel-Softmax distribution to enforce a categorical latent representation (Jang et al., 2017; Maddison et al., 2017), and employs amortised Bayesian inference (Bishop, 2006; Rezek & Roberts, 2005) to infer the model parameter distribution. This means that DyNeStE can retain the interpretability of mutually exclusive states like the HMM, while incorporating the ability to model long-range temporal dependencies using recurrent neural networks (RNNs). Just like the HMM and DyNeMo, DyNeStE is an unsupervised model in which the timing of network activations is learnt from the data without any a priori assumptions about intrinsic timescales or specific (e.g., task) timings.

In this paper, we demonstrate that DyNeStE provides an interpretable network description comparable to that of the HMM in both simulated and real MEG datasets, while also capturing long-range temporal dependencies. We first evaluate these properties using simulated data, validating DyNeStE’s ability to accurately recover ground-truth network states and underlying distributions of prolonged state activations. We then apply DyNeStE to resting-state MEG data and investigate whether it identifies a canonical set of functional brain networks that are reproducible across independent data subsets. Next, we compare DyNeStE and the HMM in their capacity to capture long-range temporal dependencies by examining whether the temporal state processes generated by each model preserve long-term dynamics of the data. Finally, to further establish DyNeStE’s reliability in MEG analysis, we assess its ability to replicate key findings from two previous HMM-based studies on memory replays (Higgins et al., 2021) and cyclical patterns of cortical network dynamics (van Es, Higgins, et al., 2025).

## 2 Methods

In this section, we describe the DyNeStE model, the datasets used, and the post-hoc analysis pipeline we employed to assess the model performance.

### 2.1 Dynamic Network States (DyNeStE)

We begin by outlining the generative model and the methodology used to infer its parameters. We then describe the training procedure of the model and choices of key hyperparameters.

#### 2.1.1 Generative Model

DyNeStE is a generative model that represents functional brain network activations through latent variables. It describes the data using a set of latent *states*, which define the time-varying statistical properties of the observed signals. The model consists of two main components: (1) an **observation model**, which parametrises each state as a distribution from which data can be sampled, and (2) a **temporal model**, which learns how these states evolve over time. Together, they form a data-generating process in which data is sampled from a distribution conditioned on the state active at each time point. An overview of the model is shown in Figure 1A, and a comparison with the HMM generative model is provided in Appendix A.1. The full mathematical formulation is presented below.

**Figure 1:**
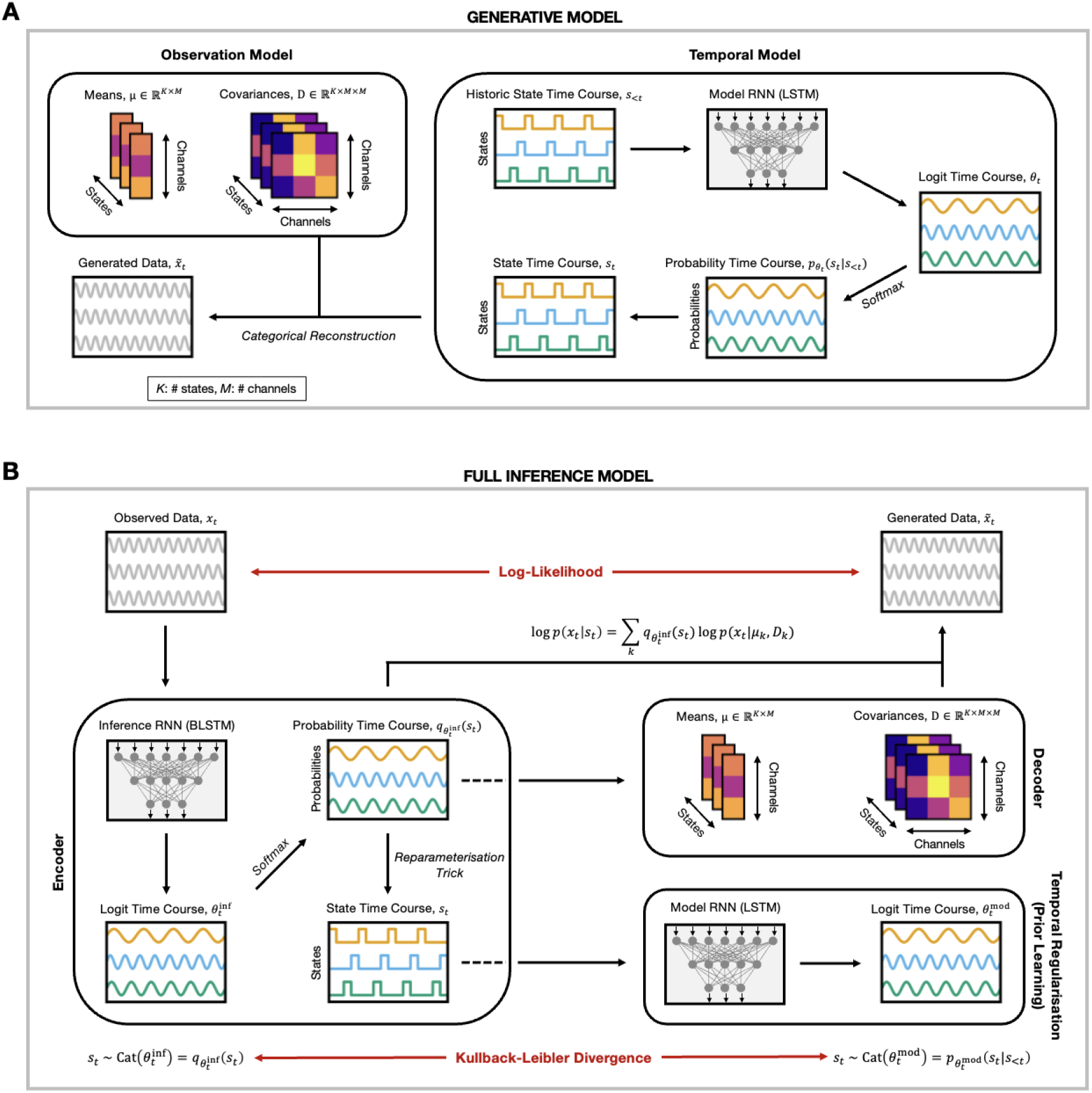
Overview of the DyNeStE model architecture. **(A)** Generative model of DyNeStE. In the temporal model, historic states *s*_*<t*_ are passed to an LSTM network, which outputs logits that parametrise a categorical distribution over the current state *s*_*t*_. The current state is then sampled from this distribution to generate the state time course. Once a state *s*_*t*_ is determined, the corresponding mean vector *µ*_*k*_ and covariance matrix *D*_*k*_ are selected from the observation model, and the generated data 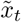 is generated from the associated multivariate Gaussian distribution. **(B)** Full inference framework of DyNeStE. The observed data *x*_*t*_ is processed by the encoder, where the inference RNN outputs logits that parametrise the variational posterior distribution. These logits are transformed via *softmax* to yield a categorical distribution over the state *s*_*t*_, from which a state is sampled using the reparametrisation trick. In the decoder, the estimated posterior is combined with state-specific means and covariances to construct the data-generating distribution *p*(*x*_*t*_|*s*_*t*_) and compute the LL. Concurrently, the sampled states are fed to the model RNN as historical input to predict the prior distribution over the next state. The KL divergence between this learned prior and the variational posterior is computed as a regularisation term. The combined LL and KL losses form the variational free energy, which is minimised through gradient descent to train the model parameters.

Let 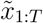 be the generated data. We have *T* as the total number of samples or time points, and 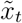 is a vector of length *M* at time *t*, where *M* is the number of channels. We assume that each data point is generated independently from a multivariate Gaussian distribution such that

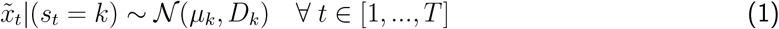

where *µ*_*k*_ ∈ ℝ^*M*^ and *D*_*k*_ ∈ ℝ^*M ×M*^ are state-specific means and covariances for each state *k* ∈ [1, …, *K*]. These mean vectors and covariance matrices comprise what we call an observation model. Because our interest lies in modelling the brain networks, for simplicity, we used the *zero-mean model* ^2^ by setting *µ*_*k*_ = 0, resulting in 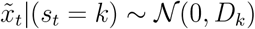.

Next, we need a temporal model that identifies which state is active at each time point. The state time course *s*_1:*T*_ is modelled as a stochastic process of state variables {*s*_*t*_}_*t*∈[1,…,*T* ]_. These variables are defined by a categorical distribution

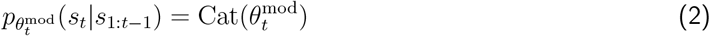

which is parametrised by the logits 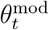 inferred using a unidirectional Long Short-Term Memory (LSTM) network (Hochreiter & Schmidhuber, 1997) called the *model RNN*. We use the model RNN to predict the future values of 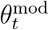 based on the historic states 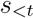3 as follows:

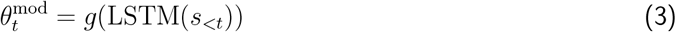

where *g* is a learnable affine transformation (i.e., a dense layer). To obtain a categorical distribution, a *softmax* function *ξ* is applied to these logits such that

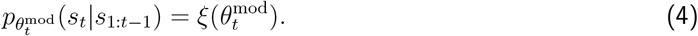

Then, 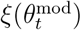 can be viewed as a vector containing the probability of each state, and *s*_*t*_ can be sampled from this probability distribution to acquire a state time course.

In summary, as a generative model, DyNeStE produces logits 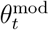 based on the history of previous states, and these logits define the current state. This state in turn determines the state-specific means and covariances from which we can generate the corresponding data points.

This formulation contrasts with that of DyNeMo (Gohil et al., 2022), which can be considered as a continuous counterpart of DyNeStE. In DyNeMo, mode^4^ probabilities are modelled by a multivariate normal distribution, with its means and covariances inferred separately as distinct logits. These mode probabilities serve as mixing coefficients, and the state-specific observation models are linearly combined according to these mixing coefficients to generate the data.

#### 2.1.2 Inference of Model Parameters

Having formulated the generative model, we now outline the inference framework for learning its model parameters, namely the logits *θ*_*t*_ and the state covariances *D*_*k*_. To do so, we structure DyNeStE as a VAE (Kingma & Welling, 2014) and use amortised variational Bayesian inference to learn a posterior distribution for the state variables *s*_*t*_ (Bishop, 2006; Rezek & Roberts, 2005).^5^ To infer the model parameters, we train DyNeStE to minimise a variational free energy loss through stochastic gradient descent using an ADAM optimiser (Kingma & Ba, 2014). A schematic overview of the entire inference framework can be found in Figure 1B.

The VAE-like architecture of DyNeStE consists of three different blocks: an encoder, a decoder, and a regularisation module which learns a prior distribution. The encoder and regularisation module constitute the temporal model of the generative process, and the decoder corresponds to the observation model.

##### Encoder

As mentioned above, adopting a VAE framework enables us to utilise amortised inference, which allows the model to process large samples of observed data and consequently scale easier (Gohil et al., 2022). Through this approach, the encoder learns the posterior distribution for each state variable and incorporates uncertainty in it.

First, given the observed data *x*_1:*T*_, we learn its hidden representation (i.e., logits) using an inference network. This hidden representation serves as a mapping from the observed data to the parameters of the variational posterior distribution. Here, we call an inference network the *inference RNN*, which is a bidirectional LSTM (BLSTM) network. Hence, we have

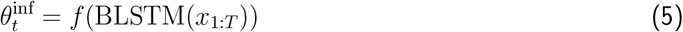

where 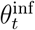 are the logits of the inference RNN, and *f* is a learnable affine transformation.

Next, the resulting logits are passed through a softmax function to calculate the state probabilities *α*_*t*_ such that

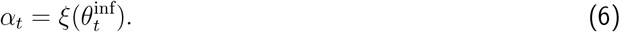

Because we are modelling states, these probabilities can be taken as a categorical posterior distribution 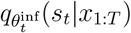 at each time point. However, we cannot directly sample a state *s*_*t*_ from this categorical distribution because gradients cannot propagate through categorical variables (i.e., one-hot vectors). Sampling from a categorical distribution is a non-differentiable operation due to its discrete and non-smooth nature. To circumvent this problem, we apply a *reparametrisation trick* using the *Gumbel-Softmax* distribution (Jang et al., 2017; Maddison et al., 2017). This approach enables differentiable approximation of categorical samples by transforming them into continuous representations. Specifically, we approximate a posterior distribution as

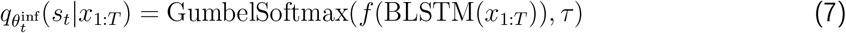

where *τ* is the temperature of the *softmax* function. In practice, we anneal *τ* from 1 to 0.06 to stabilise the learning process, enforcing the model to begin by learning a smooth, continuous distribution and gradually transition into a more discrete, categorical distribution. As a result, we can sample our discrete random variable *s*_*t*_ from a variational posterior, allowing us to introduce stochasticity and capture the uncertainty in *s*_*t*_ during model training.

##### Decoder

In the decoder block, the likelihood of generating the observed data is estimated given the sequence of states:

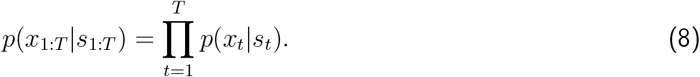

Here, we factorise the likelihood, assuming that the data at each time point is independent and only depends on the state vector at the corresponding time point. We further assume that the likelihood function is a multivariate normal distribution such that

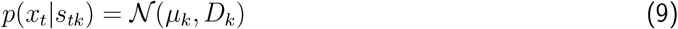

where *s*_*tk*_ indicates whether a state *k* is active or not. As in Section 2.1.1, we use a zero-mean model and set *µ*_*k*_ = 0.

We learn the state-specific covariances as trainable free parameters (i.e., point estimates). Because the covariance^6^ is required to be positive semi-definite, we parametrise *D*_*k*_ using the Cholesky decomposition as

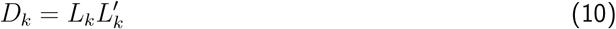

where *L* is a lower triangular matrix called a Cholesky factor and ^*′*^ denotes a matrix transpose. We add a small positive value to the diagonal of the Cholesky factor to enhance training stability and make the covariance matrix strictly positive definite. This addition can be thought of as a form of the Tikhonov regularisation that enhances numerical stability of our solution.

##### Temporal Regularisation (Prior)

In contrast to a VAE, which has a fixed prior (i.e., the same prior for all time points), we include temporal regularisation by learning a prior that predicts *s*_*t*_ based on the previous state history *s*_*<t*_. As outlined in Section 2.1.1, the prior

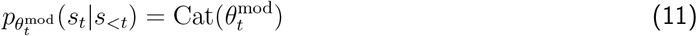

is modelled as a categorical distribution parametrised by a logit vector 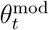, which is inferred using the model RNN (Equation (3)). The model RNN takes as input the previous states *s*_*<t*_, sampled from the posterior distribution of the encoder. By measuring the Kullback-Leibler (KL) divergence (Kullback & Leibler, 1951) between the variational posterior and the prior, the learned posterior can be regularised to remain close to the prior distribution and incorporate the influence of previous states (see Section 2.1.3).

#### 2.1.3 Training DyNeStE

With the subcomponents of the DyNeStE model architecture outlined, the next step is to understand how they are combined as a whole model and trained to minimise the loss function. For DyNeStE, the loss ℒ is given by the variational free energy

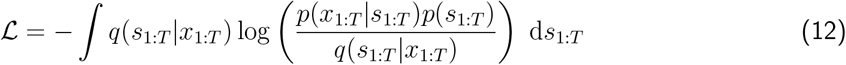

where *q*(*s*_1:*T*_ |*x*_1:*T*_ ) is the posterior, *p*(*s*_1:*T*_ ) is the prior, and *p*(*x*_1:*T*_ |*s*_1:*T*_ ) is the likelihood. This loss can be divided into two separate terms – negative log likelihood (LL) and KL divergence:

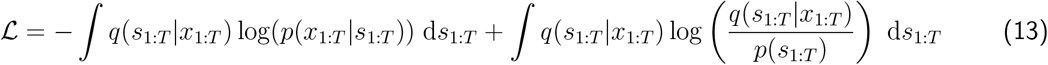

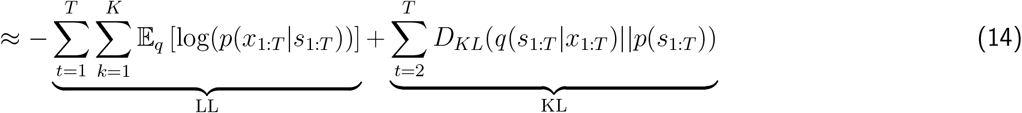

where 𝔼(·) is the expectation, and *D*_*KL*_(·||·) is the KL divergence.^7^ For the complete mathematical derivation of the loss function, readers should refer to Appendix A.2, where we delineate how the individual model components are integrated together. The appendix also elaborates on how the variational posterior, likelihood, and prior (defined in Equations (7), (9), and (11), respectively) are combined to yield Equation (12).

The three components of DyNeStE are trained jointly in an end-to-end fashion. In brief, the observed data is processed through the encoder to infer the variational posterior distribution. This posterior is then combined with the log-likelihood function from the decoder to define the data-generating process *p*(*x*_*t*_|*s*_*t*_), yielding a negative LL loss that quantifies the reconstruction error between the observed and generated data.

Concurrently, states sampled from the inferred posterior distribution serve as a historical input to the regularisation module, which learns a prior distribution *p*(*s*_*t*_|*s*_*<t*_). Here, the KL divergence between the learned posterior and prior is computed as a regularisation loss term.

Finally, these two components are added up to form the objective function in Equation (14), which is minimised through optimising trainable parameters of DyNeStE. Several additional training techniques were employed to enhance optimisation and model convergence (see Appendix A.3 for details).

In terms of training and inference, DyNeMo shares the same overall structure as DyNeStE (i.e., the encoder, decoder, and temporal regularisation) but differs in how the variational posterior and prior are defined. In DyNeMo, both are modelled as continuous multivariate normal distributions, which modify the KL divergence term in the objective function. As a result, the modes are represented in a continuous latent space, and the log-likelihood loss is evaluated under a multivariate normal likelihood, weighted by the variational posterior through Monte Carlo estimation (see SI 1.1 in Gohil et al., 2022).

### 2.2 Datasets

In this study, three datasets were used to train and evaluate DyNeStE and the HMM: one simulated dataset to validate DyNeStE’s modelling and inference capabilities, and two resting-state MEG datasets to assess its applicability and effectiveness in MEG analysis compared to the HMM.

#### 2.2.1 Simulation

For the simulation data, we generated a multivariate time series with 3 hidden states, 11 channels, and 256,000 samples using an HSMM (Yu, 2010), whose observation model was a multivariate Gaussian distribution with a zero mean vector and real data-driven covariance matrix for each state. Unlike standard HMMs, the HSMM explicitly models state lifetimes (i.e., the duration for which a state remains active), allowing us to probe whether DyNeStE can capture long-range temporal dependencies in its learnt parameters.

To make long-lived states more probable, we defined the lifetime distribution as a gamma distribution with shape and scale parameters of 10 and 5, respectively. State lifetimes were sampled from this distribution, with the sequence of states determined by a transition probability matrix

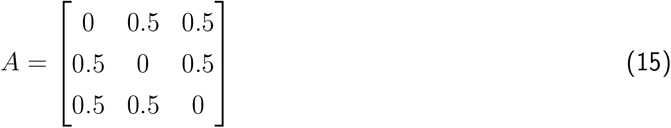

in which self-transitions are excluded, as state lifetimes were already specified, and the probability of transitioning to any other state is uniform. The ground-truth state time course and state covariances are shown in Figure 2A and B.

**Figure 2:**
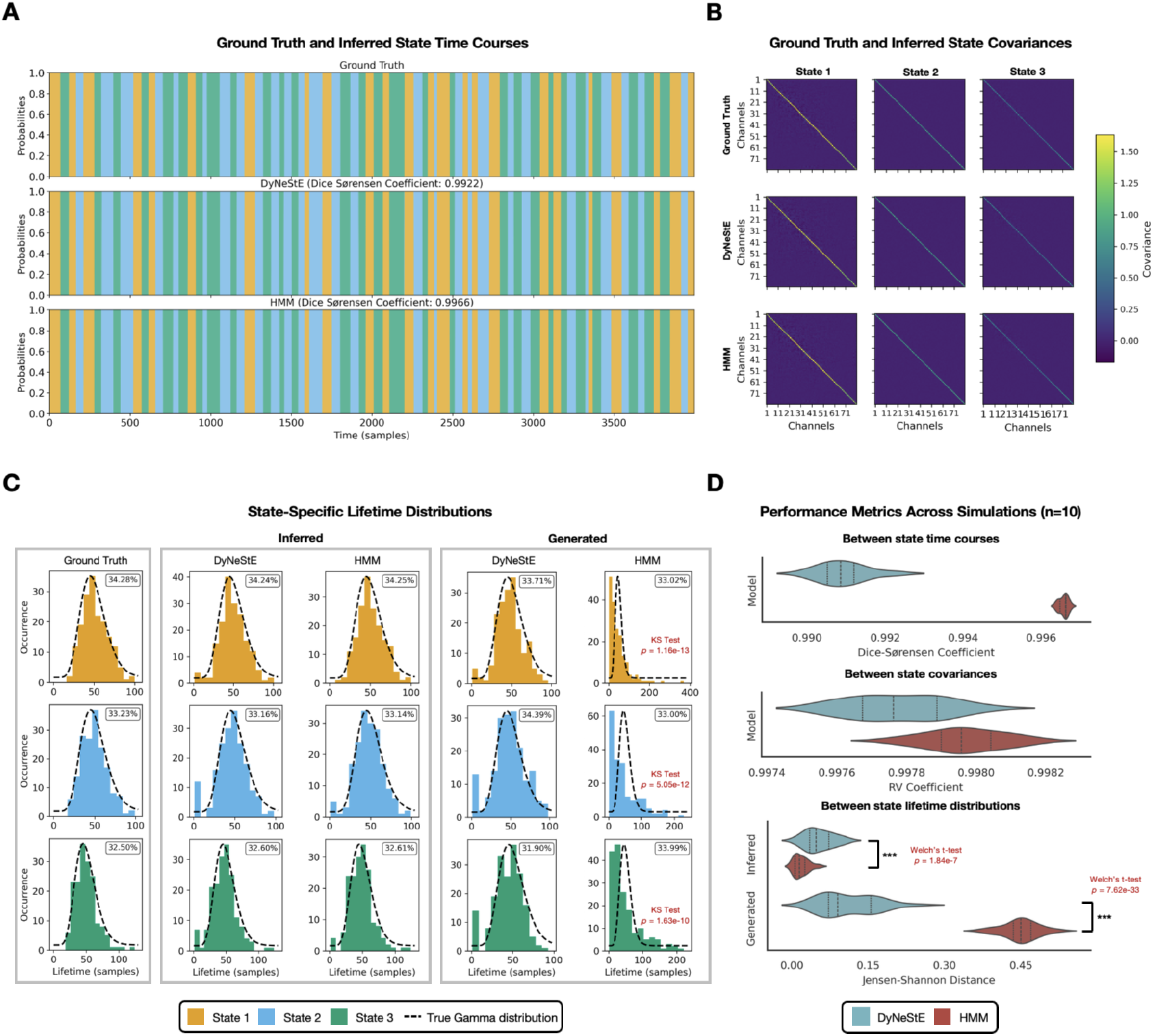
Validation of DyNeStE using simulated data. **(A)** State time courses inferred by DyNeStE and the HMM and their ground truth are shown for the first 4,000 samples, with different states shown as different colours (see legend in panel C). For each model, the Dice-Sørensen coefficient between the ground-truth and inferred state time courses is reported. **(B)** State covariance matrices inferred by DyNeStE and the HMM are shown alongside the ground truth. **(C)** Distributions of state lifetimes are shown for the state time courses that were inferred from the data, generated by each inferred model, and used in the data simulation (i.e., the ground truth). The ground-truth gamma distribution used to generate the simulated data is also overlaid as a black dotted line. Insets display the fractional occupancy of each state. The KS test was used to compare inferred and generated state lifetime distributions against the ground truth; only significant results are highlighted in red. **(D)** Across ten simulation runs, performance metrics of each model are shown as violin plots: Dice-Sørensen coefficients between the ground-truth and inferred state time courses (top), RV coefficients between the groundtruth and inferred state covariances (middle), and JS distances between the ground-truth and inferred/generated lifetime distributions (bottom). For comparisons of the JS distance between the two models, the Welch’s t-test was applied to both inferred and generated lifetime distributions. P-values are reported in red (***: *p <* 0.001, **: *p <* 0.01, *: *p <* 0.05, n.s.: non-significant).

#### 2.2.2 Real MEG Data

For the real MEG data, we used two datasets in this paper:

- **Nottingham MEGUK Dataset** consists of resting-state, eyes-open MEG data collected using a 275-channel CTF scanner at a sampling frequency of 1.2 kHz. It contains scans from 65 healthy subjects, each lasting approximately 5 minutes (see Appendix A.4 for the subject demographics). This dataset was collected at the University of Nottingham as part of the UK MEG Partnership.
- **Replay Dataset** was introduced in Liu et al. (2019) and further analysed in Higgins et al. (2021). Of the three types of datasets described in Higgins et al. (2021), we used “Dataset A”, which we refer to as the Replay dataset in this paper. The resting-state MEG scans were acquired using a 275-channel CTF scanner at a sampling frequency of 600 Hz. These data were collected from 21 healthy subjects, with two sessions per subject. Each recording was approximately 10 minutes long.

Detailed descriptions of these datasets, along with the preprocessing and source reconstruction procedures we applied, can be found in Gohil, Huang, et al. (2024), Higgins et al. (2021), and Huang et al. (2025). For completeness, we briefly summarise the preprocessing and source reconstruction pipeline below. The entire pipeline was implemented using the osl-ephys toolbox (van Es, Gohil, et al., 2025).

##### Data Preprocessing

For the Nottingham MEGUK dataset, the raw time-series data were first bandpass filtered between 0.5 and 125 Hz and notch filtered at 50 Hz and 100 Hz. The data were then downsampled to 250 Hz. Abnormally noisy segments and channels in the data were automatically detected using the generalised extreme Studentised deviate algorithm (Rosner, 1983). Next, sensor data were decomposed into 64 components using FastICA (Hyvarinen, 1999), and components showing a high correlation with electrooculogram (EOG) and electrocardiogram (ECG) were marked as noise and removed. To keep the data dimension consistent across subjects, any sensor channels identified as bad in earlier steps were interpolated from ICA-cleaned data via the spherical spline interpolation method (Perrin et al., 2019).

For the Replay dataset, the raw time-series data were bandpass filtered from 1 to 45 Hz and subsequently downsampled to 250 Hz. The recordings were visually inspected to remove any noisy segments or channels. FastICA (Hyvarinen, 1999) was utilised to decompose the sensor data into 150 components. Artefact components were identified based on their spatial topography, time courses, kurtosis, and frequency spectrum and were removed from the data.

##### Source Reconstruction

For the Nottingham MEGUK dataset, the preprocessed data were coregistered with each subject’s structural MRI (sMRI) scans and digitised headshape and fiducial points acquired with a Polhemus pen. Following coregistration, the data were bandpass filtered between 1 and 45 Hz and reconstructed onto an 8-mm isotropic grid using a linearly constrained minimum variance (LCMV) beamformer (Van Veen & Buckley, 1988).

Our use of the LCMV beamformer is consistent with prior studies concerning the TDE-HMM (Vidaurre et al., 2018) and DyNeMo (Gohil et al., 2022), which allows for direct comparability with these established frameworks. LCMV has been shown to perform well in suppressing noise, detecting point-like sources, and estimating coherence compared to the Minimum Norm Estimate (MNE) approach (Hincapié et al., 2017). At the same time, it has also been argued that no single method is superior among commonly used approaches such as LCMV, DICS (Dynamic Imaging of Coherent Sources), MNE, and eLORETA (exact Low-Resolution Electromagnetic Tomography), as each entails distinct strengths and limitations; consequently, their complementary use has been proposed as an alternative strategy (Halder et al., 2019). While we do not expect our choice of source reconstruction method to substantially affect the conclusions we draw here, it is important to note that different techniques may be more favourable depending on the specific scientific hypothesis under investigation.

The resulting voxels were parcellated into 38 anatomically defined regions using the Giles parcellation (Colclough et al., 2016). Source reconstructed signals were extracted using the principal component analysis (PCA)^8^, with each parcel time course defined as the first principal component across all voxels constituting the corresponding parcel. To minimise spurious correlations between parcels and reduce source leakage, we applied the symmetric multivariate leakage reduction technique (Colclough et al., 2015), which removed all zero-lag correlation between parcel time courses.

The use of LCMV beamformer and PCA results in parcel time courses with arbitrarily oriented dipole signs across subjects, which obscures group-level analysis. To correct this, a greedy search-based random sign-flipping algorithm was used for alignment.

The Replay dataset followed the same source reconstruction pipeline, except that coregistration relied solely on fiducial markers due to the absence of sMRI scans^9^. All subsequent analyses were performed on these parcel-level time courses.

#### 2.2.3 Data Preparation for Model Training

For model training, a few additional data preparation steps were applied to parcel-level time courses of both simulated and real datasets. First, the data were time-delay embedded with *±*7 lags to enable a model to infer states with distinct spatiotemporal patterns as part of its observation model (Gohil et al., 2022). Next, we used PCA to reduce the dimensionality of the embedded data to 80 channels, thereby reducing computational memory requirements. Lastly, to enhance optimisation stability and facilitate convergence, the transformed data were standardised over time (Géron, 2022), with standardisation applied independently to each subject or session.^10^

### 2.3 Model Training

We trained DyNeStE and the TDE-HMM^11^ (Vidaurre et al., 2018) on the simulation and Nottingham MEGUK datasets. The prepared data (in Section 2.2.3) served as an input for model training. For the real dataset, data from all subjects/sessions were concatenated. This means that the inferred brain networks are shared across recordings, providing a group-level description of brain activity. The input data were shuffled and batched prior to training.

Each model was trained 10 times, and the run with the lowest variational free energy was selected for post-hoc analysis (see Appendix A.3.2 for the training loss curves). The remaining runs were used for statistical analysis. Because state ordering is arbitrary across runs and model types, we used the Hungarian algorithm to match the orders based on the state covariance matrices, followed by manual visual alignments if necessary. The model hyperparameters for each dataset are listed in Appendix A.3.3.

After training, the inferred state probabilities (see Equation (6) for DyNeStE) and observation model parameters of each model were saved for the post-hoc analysis. State probabilities were converted to a state time course by taking the argmax at each time point (i.e., the point-wise maximum a posteriori (MAP) estimate). We refer to this as the **inferred state time course**.

For the Replay dataset, no model was trained; rather, we applied models pre-trained on the Nottingham MEGUK data to infer state time courses. This approach allowed us to avoid potential confounds from training on data that were source reconstructed without sMRI, while also enabling direct replication of previous findings in Higgins et al. (2021).

### 2.4 Data Generation from DyNeStE and HMM

To assess how well the models capture long-term dependencies, we generated synthetic data using the trained DyNeStE and HMM (see Appendix A.1) and compared the temporal structure of these data to that of the observed data (described in Section 2.9).

For DyNeStE, we first sampled an initial state from a Gumbel-Softmax distribution using the final annealed temperature. This initial state was then fed into the learnt model RNN to predict the next state. This process was repeated over time, with each predicted state fed back into the RNN to produce a continuous sequence of states. At each time step, we applied a *softmax* function to the model RNN logits to obtain the state probabilities *α*_*t*_, on which an argmax operation was subsequently applied to acquire a state time course. For HMM, state time courses were generated by simulating the HMM using the learnt transition probability matrix.

The resulting outputs of both models are referred to as **generated state time courses**. For the simulation, we generated data at 10% of the full dataset size (i.e., 25,600 samples^12^) to reduce the computational cost; for the Nottingham MEGUK dataset, state time courses were generated for 5 minutes for each of the 65 subjects.

### 2.5 Simulation Analysis

From the inferred (or generated) state time courses, we can extract the duration of each state activation as a state lifetime and construct a lifetime distribution. This distribution was compared against the ground truth using a two-sample Kolmogorov-Smirnov test with an alpha threshold of 0.05. Multiple comparisons were addressed with Bonferroni correction for 12 tests, covering three states, two models, and both inferred and generated state time courses.

To confirm consistency in the model performance, DyNeStE and the HMM were each trained on 10 independent simulation runs. For each run, we calculated the Dice-Sørensen coefficient between groundtruth and inferred state time courses, as well as the RV coefficient between ground-truth and inferred state covariances. The model’s ability to consistently recover lifetime distributions was measured using the Jensen-Shannon (JS) distance between ground-truth and inferred/generated lifetime distributions. We compared the JS distances between DyNeStE and the HMM using a Welch’s t-test, after verifying the assumptions of normality and equal variance.

### 2.6 Power and Functional Connectivity

After training DyNeStE and the HMM on the real data, the inferred states were interpreted as dynamic resting-state networks (RSNs), and their network profiles were characterised using power spectral densities (PSDs), power maps, and functional connectivity (FC) networks.

For each state, subject-level PSDs and cross-spectral densities (CSDs) were estimated over the frequency range of 1–45 Hz using the multi-taper method with seven tapers and a time-half bandwidth of 4. Parcel time courses were z-score standardised prior to PSD computation, and data segments corresponding to each state’s activation periods were extracted based on the inferred state time courses. Subject-level spatial power maps were computed by averaging PSDs across the frequency range of 1.5–20 Hz.

FC was quantified using coherence measures. To calculate subject-level coherence networks, we first computed the coherency spectrum from the PSDs and CSDs and then assigned the mean coherence value within the 1.5–20 Hz frequency band as the edge weight for each parcel pair.

We chose coherence as our FC measure because both DyNeStE and the HMM are formulated to infer coherence-based network states from CSDs estimated from TDE data (Gohil et al., 2022). Because time-delay embedding preserves the cross-spectral structure of neural time series by design, coherence becomes a natural choice for quantifying frequency-specific oscillatory phase synchronisation between brain regions. As a phase-based metric, it is also well suited to dynamic analyses in which transient, time-lagged inter-regional interactions are of interest. Consistent with this rationale, coherence has been frequently used in prior dynamic network studies employing similar modelling frameworks (e.g., Vidaurre et al., 2016; Vidaurre et al., 2018; Gohil, Huang, et al., 2024).^13^

Group-level PSDs, power maps, and FC networks were obtained by averaging across all subjects, with each subject-wise feature weighted by its respective data length. For visualisation purposes, power maps and FC networks were demeaned by subtracting their mean across all states, weighted by the duration of state activations. Additionally, FC network edges were thresholded at the 95th percentile.

### 2.7 Split-half Reproducibility

To determine whether the dynamic RSNs of DyNeStE are reproducible, we split the Nottingham MEGUK dataset into two halves and independently inferred networks from each half. Following the procedure in Section 2.3, DyNeStE was trained 10 times on each subset, and the best-performing runs were selected for comparison. We first computed power maps from the two halves and compared them qualitatively. To quantify the reproducibility across the split-halves, we then calculated cosine similarities between corresponding state-wise power maps and assessed their statistical significance using a maximum-statistic permutation test (see Appendix D in Huang et al. (2025) for details).

### 2.8 Network Dynamics

Next, we employed three distinct metrics to compare the network dynamics of state transitions inferred by DyNeStE and the HMM.

#### State time course correlation

For each subject, we computed pairwise Pearson correlations between inferred state time courses of the two models. These values were then averaged across subjects to obtain group-level, state-wise correlation measures.

#### Riemannian distance

Using group-level covariances from the inferred observation models, we measured spatio-spectral differences between the two models by calculating the Riemannian distances between their respective covariance matrices.

#### Summary statistics

Four summary metrics were adopted to quantify the temporal characteristics of the RSNs (Baker et al., 2014; Cho et al., 2024). These metrics were calculated from the inferred state time courses for each subject and include:

- Fractional occupancy: the proportion of total time spent in each state
- Mean lifetime: the average duration of each state activation before transitioning into another state
- Mean interval: the average time elapsed between successive visits to the same state
- State switching rate: the frequency of state activations per unit time (second)

### 2.9 Measures of Long-Range Temporal Dependencies

The long-range temporal dependencies were measured from the inferred and generated state time courses using two metrics that recapitulate different aspects of the temporal structure.

#### 2.9.1 Fano Factor

First, we computed the Fano factor from state time courses at the subject/session level across multiple window lengths (Higgins et al., 2021). For each window length, we segmented the time course into non-overlapping windows and counted the number of state activations for each state within each window. We then calculated the mean *µ* and variance *σ*^2^ of activation counts across all windows. The Fano factor for a window length *L* was therefore defined as

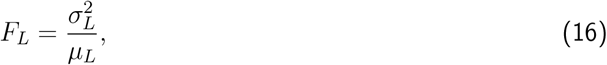

providing a measure of temporal variability in state activation patterns.

Here, the Fano factor quantifies whether state activations are clustered or regularly distributed over time. Values equal to 1 indicate memoryless Poisson-like processes with regular, independent events. On the other hand, values greater than 1 reflect irregular state activations that suggest the presence of long-term dependencies. Such temporal clustering implies that the probability of future activations may depend on the history of past events, which indicates a stronger long-range temporal structure.

#### 2.9.2 Time-Lagged Mutual Information

To quantify statistical dependencies between state activations, we also calculated the mutual information (MI) between state time courses across different time lags. MI was computed pairwise for all unique state combinations at each lag, with negative and positive lags representing backward and forward temporal shifts, respectively. Because of the variability in entropy values across state pairs, MI was normalised by the geometric mean of the marginal entropies of the two states (in nats). For each state, MI values were averaged over all state pairs, excluding the self-information terms. This yielded state-averaged MI profiles across lags for each subject, providing a time-resolved measure of inter-state dependencies.

When MI decays slowly over longer time lags, it suggests that the activation of one state carries information about past and future state activations over extended time horizons. In contrast, a rapid drop in MI to near-zero values at longer lags indicates predominantly short-range dependencies. Sustained MI at long lags thus implies that the state time courses retain memory of past configurations and that the model captures long-range temporal dependencies.

#### 2.9.3 Statistical Analysis

Finally, to quantitatively compare the ability of DyNeStE and the HMM to capture long-range temporal dependencies, we tested whether the Fano factor and MI calculated from each model’s inferred (or generated) state time courses differ significantly. For each state, statistical significance was assessed using a max-t cluster permutation test (as implemented in MNE-Python software v1.9.0; Gramfort et al., 2013) comparing DyNeStE and the HMM across subjects, with clusters defined over windows/lags. This test controls for multiple comparisons across windows/lags by evaluating cluster-level statistics against a max-t distribution. To further account for testing across multiple states, the resulting p-values were adjusted using a Bonferroni correction over the number of states.

### 2.10 TINDA Analysis

Another way to examine the model’s ability to capture long-range temporal dependencies is by looking at whether the cyclical structure known to be present in the activations of dynamic RSNs are learnt by the models. To extract this cyclic pattern from the state time courses, we used a method called the Temporal Interval Network Density Analysis (TINDA), introduced in (van Es, Higgins, et al., 2025). The methods described below are adapted from this study, and readers should refer to the original publication for comprehensive details. We provide a brief overview here for clarity.

#### 2.10.1 Fractional Occupancy Asymmetry Matrix

Conceptually, TINDA measures the tendency of one state to precede or follow the other states and gauges the presence of cyclic patterns in the state dynamics. To do so, it quantifies directional asymmetries in network transitions over inter-state intervals (ISIs) of varying lengths. For each reference state, all intervals between consecutive occurrences of that state are identified and divided into equal first and second halves. For every other network state, we compute its fractional occupancy (FO) separately in the first and second halves of the intervals. The FO asymmetry for a given state pair is defined as the difference in FO between the two halves, averaged over all ISIs. Repeating this for all pairs of reference and target states yields the FO asymmetry matrix of size *K × K*, whose entries reflect the magnitude and direction of temporal bias between the states. These FO asymmetry matrices were computed for each subject.

#### 2.10.2 Detection of Cyclical Structure and Cycle Strength

The FO asymmetry matrix can be conceived as a weighted directed graph, where nodes are network states and edges are the strength of asymmetries between state pairs. To determine whether these pairwise asymmetries form a coherent cycle, we positioned states onto a unit circle and searched for the ordering that maximised the net clockwise transitions. Specifically, for each pair of states, we projected the corresponding FO asymmetry value tangentially to the unit circle. We then defined *cycle strength* as the normalised sum of these tangential projections, with a value of +1, 0, and −1 representing a perfectly clockwise cycle, no net directionality, and a perfectly counterclockwise cycle, respectively. The optimal cycle structure was identified as the one that maximised this cycle strength over permutations of state positions on the unit circle. The final cycle strengths from the inferred and generate state time courses were compared between DyNeStE and the HMM using two-sample *t*-test (after verifying the normality and equal variance assumptions) with an alpha threshold of 0.05.

#### 2.10.3 Circle Plot Visualisation and Timescale-specific Analysis

Once the optimal ordering of states was established, we generated a circular plot in which each state was positioned at its corresponding phase angle. Directed edges were drawn between state pairs whose FO asymmetries were strongest (i.e., statistically significant). The statistical significance was tested at the group level using a two-tailed paired-samples *t*-test for each directed connection, with the alpha threshold of 0.05 Bonferroni-corrected for *K*^2^ − *K* = 132 comparisons.

To examine potential variation in cyclical structure across temporal scales, the entire TINDA analysis was repeated after dividing the distribution of ISIs into five equally populated bins (i.e., quintiles). FO asymmetry estimation and cycle detection were applied separately within each bin, enabling identification of the timescales at which cyclical dynamics were most pronounced. For each quintile, the cycle strengths were compared between DyNeStE and the HMM using Welch’s *t*-test (after verifying the normality and equal variance assumptions), with an alpha threshold of 0.05 and Bonferroni correction for multiple comparisons across quintiles.

### 2.11 Replay Data Analysis

In addition to the TINDA analysis, we assessed whether DyNeStE can replicate the findings from the previous HMM-based study (Higgins et al., 2021) by investigating the network dynamics linked to memory replays. The methods described below are adapted from this study, and readers should refer to the original publication for comprehensive details. We provide a brief overview here for clarity.

For the Replay data analysis, we first detected replay events following Liu et al. (2019). In particular, multivariate classifiers were trained on the functional localiser data to find task-evoked MEG patterns associated with each learnt stimulus. They were then applied to resting-state data to identify rapid sequential reactivations of these patterns in the task-defined order. The resulting probabilities of these reactivations were thresholded at the 99th percentile to obtain replay event times. RSNs and their state time courses were inferred from the data as described in Section 2.3.

Using the replay events and state (probability) time courses, we computed four metrics:

#### Replay-evoked Network Activations

Replay-evoked network activations were quantified as time-locked changes in state probabilities around replay onsets. For each session, the sequence of state probabilities (*α*_*t*_) was baseline-corrected by subtracting the session-specific mean across all time points. Then, the state probability time courses were epoched into windows spanning 0.5 s before and after each replay event. Within each session, the statewise, event-locked probability traces were averaged over all corresponding epochs to get a session-level replay-evoked network activation.

#### Mean Replay Intervals Given Active RSN States

Next, we identified the RSN state active at each replay onset and computed the interval to the subsequent replay event. For each state, these inter-replay intervals were aggregated across both sessions per subject, and the mean interval time was computed over all replay intervals. Subjects with fewer than 10 replay events for a given state were excluded to prevent high-variance estimates.

#### Mean Replay Rates Given Active RSN States

Replay rates were calculated as the number of replay events occurring during activation periods of each RSN state, divided by the total duration of that state activation. For each state, these measures were aggregated across both sessions per subject, and the mean replay rates were computed over all activation instances.

#### Fano Factor of RSN States

As described in Section 2.9.1, Fano factors of the RSN states were computed as a metric of long-term dependencies and compared between DyNeStE and the HMM. The effect size was defined as the mean group difference, with variance estimated as the standard error of this mean group difference.

#### 2.11.1 Statistical Analysis

The statistical significance of the four metrics was assessed using non-parametric tests. For the replay-evoked network activations, we applied a one-sample max-t cluster permutation test over the time dimension to identify time points at which these state probabilities are significantly greater than the baseline value. For mean replay intervals and replay rates, we used a one-sample max-t permutation test, comparing each state’s replay intervals or rates against the subject-averaged mean values computed across all other states. For the Fano factor, we applied a max-t cluster permutation test over the time dimension to identify differences between DyNeStE and the HMM. These tests were performed separately for each state using MNE-Python v.1.9.0 (Gramfort et al., 2013), and resulting p-values were Bonferroni-corrected for multiple comparisons over the number of states.

## 3 Results

### 3.1 Simulation Analysis

To validate the design choices underlying DyNeStE and test its efficacy, we first simulated data using the HSMM (see Section 2.2.1) and investigated whether DyNeStE can reliably infer categorical latent states and capture long-range temporal dependencies. This validation aims to confirm whether DyNeStE can correctly identify a known latent description of the data as a latent variable model in a simple case; evidence of robustness under real-world conditions is presented in Sections 3.2 and 3.3.

#### 3.1.1 DyNeStE learns categorical latent states in simulated data

We inspected the states inferred by DyNeStE by examining the learnt model parameters of its temporal and observation model (Fig. 1), i.e., the state time courses^14^ and state covariance matrices. DyNeStE correctly inferred state transitions in a categorical and discrete manner, comparable to the HMM, achieving a Dice-Sørensen coefficient greater than 0.99 between the inferred and ground-truth state time courses (Fig. 2A). Moreover, its observation model was qualitatively identical to that of the HMM, with both models accurately recovering the ground-truth multivariate normal distribution of the simulated states (Fig. 2B).

#### 3.1.2 DyNeStE captures long-range temporal dependencies in simulated data

Having established that DyNeStE can learn categorical network states, we next verified its ability to capture long-range temporal dependencies. As the simulated data were generated with states persisting over extended periods, whose state lifetimes were governed by the pre-defined gamma distributions, we examined whether DyNeStE could recover these true state lifetime distributions.

Both DyNeStE and the HMM successfully inferred existing long-range temporal dependencies, matching the ground-truth state lifetime distributions (Fig. 2C, middle). However, this agreement could arise trivially from data-driven inference, rather than the generative models themselves capturing long-term network dynamics in the model parameters that describe such temporal dynamics (i.e., the transition probability matrix in the HMM and the model RNN in DyNeStE). To test this, we sampled state time courses from the trained DyNeStE and HMM models and evaluated their lifetime distributions. While the data generated by DyNeStE reproduced the ground-truth distributions, HMM-generated data yielded exponential lifetime distributions for all states, suggesting that it can only capture short-range dependencies, as is expected given its first-order Markovian constraint (Fig. 2C, right). These results indicated that the recurrent modelling of network states by DyNeStE can encode the relationship between state activations over long periods of time within its parameters, whereas the HMM cannot.

We further assessed if the results above are robust across multiple simulation runs (Fig. 2D). Both DyNeStE and the HMM consistently achieved the Dice-Sørensen coefficients greater than 0.99 between inferred and ground-truth state time courses, as well as RV coefficients above 0.9 between inferred and ground-truth covariance matrices. Consistency in capturing long-range temporal dependencies was quantified with JS distances between the ground-truth and inferred/generated lifetime distributions. While both models produced inferred distributions nearly identical to the ground truth across the runs, a significant difference between DyNeStE and the HMM (Welch’s t-test; *p* = 1.84 *×* 10^−7^, *t* = 6.19) was observed, largely due to DyNeStE exhibiting a greater prevalence of transient state activations (see Section 4.5.1). For the generated lifetime distributions, there was a clear distinction in which the HMM deviated substantially from the ground truth, while DyNeStE matched it with greater precision (Welch’s t-test; *p* = 7.62 *×* 10^−33^, *t* = −29.3).

### 3.2 Real Data Analysis: Nottingham MEGUK

#### 3.2.1 DyNeStE infers plausible categorical dynamic brain networks

Following validation on the simulated data, we applied DyNeStE to resting-state MEG recordings to assess its model performance on real data. We trained DyNeStE on the Nottingham MEGUK dataset, from which 12 dynamic functional brain networks were inferred (Fig. 3). Qualitatively, these RSNs aligned well with canonical brain networks reported in previous HMM-based studies (Cho et al., 2024; Gohil et al., 2022), including the motor (State 1), visual (State 5), anterior default mode (State 7), posterior default mode (State 9), and temporal networks (States 6 and 8). They also aligned well with the states inferred by the HMM when trained on the same dataset (Fig. A5). Altogether, these results demonstrated that DyNeStE can reliably recover plausible RSNs in a discrete, categorical manner.

**Figure 3:**
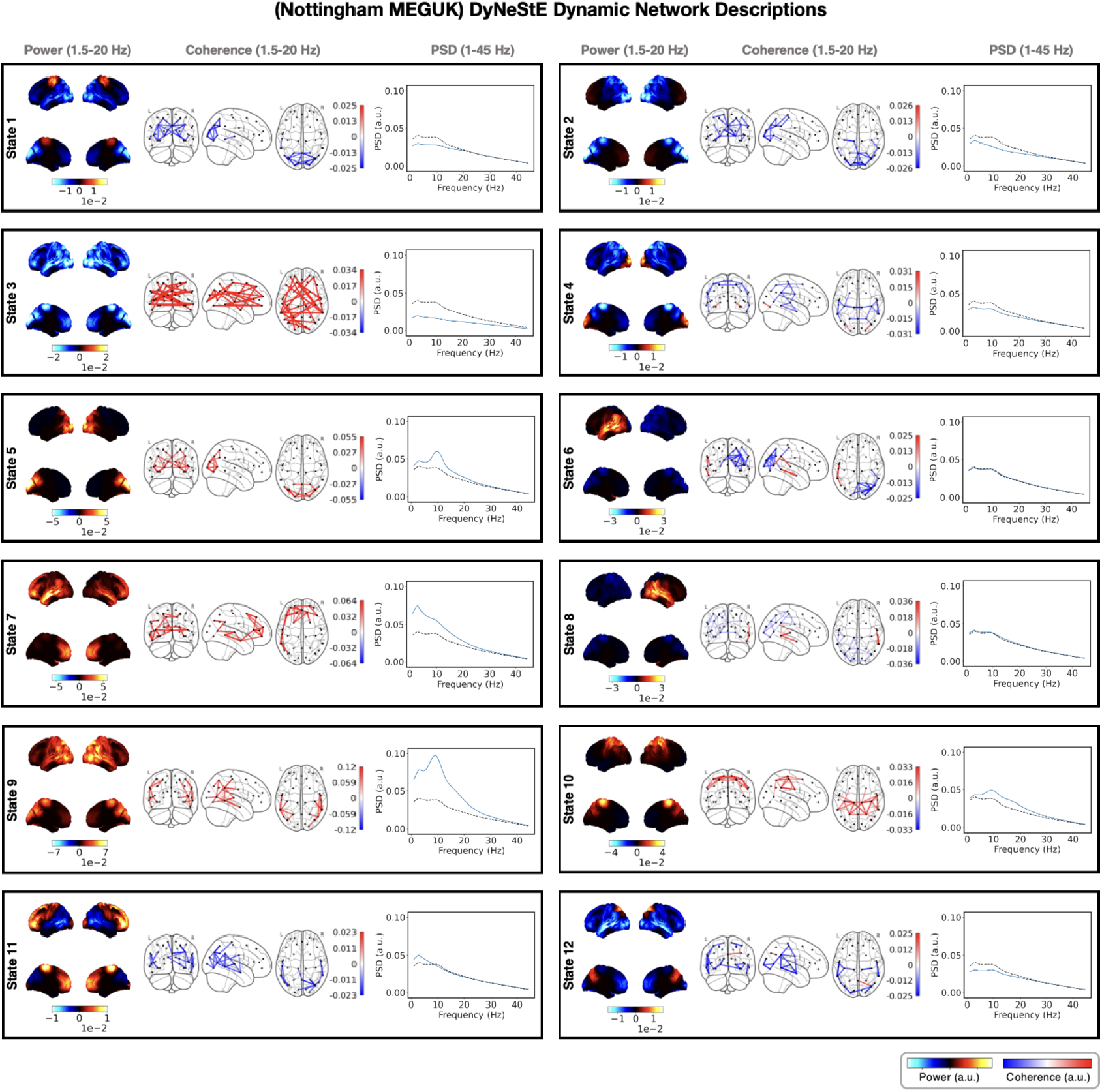
Dynamic resting-state MEG networks inferred by DyNeStE. Twelve states were identified by DyNeStE using resting-state MEG recordings of 65 subjects in the Nottingham MEGUK dataset. Each panel summarises network descriptions of one state, namely the group-level power map (left), FC network (middle), and parcel-averaged PSD (right). The power maps display lateral and medial cortical surfaces at the top and bottom, respectively. The FC networks illustrate edges with the top 3% coherence values (regardless of sign). Both power maps and FC networks are shown relative to their average across all states. The PSD of each state (blue) is plotted alongside the state-averaged PSD (black dotted line).

The dynamics of the RSNs further supported the categorical nature of DyNeStE states. The state probability time courses estimated by DyNeStE were nearly one-hot, producing mutually exclusive state time courses that closely resembled those inferred by the HMM (Fig. 4A). Such similarity between the two models was confirmed quantitatively. The state time courses of both models were highly correlated across all states (Fig. 4B), and their observation models described similar state distributions, as demonstrated by short Riemannian distances between their state covariance matrices (Fig. 4C).

**Figure 4:**
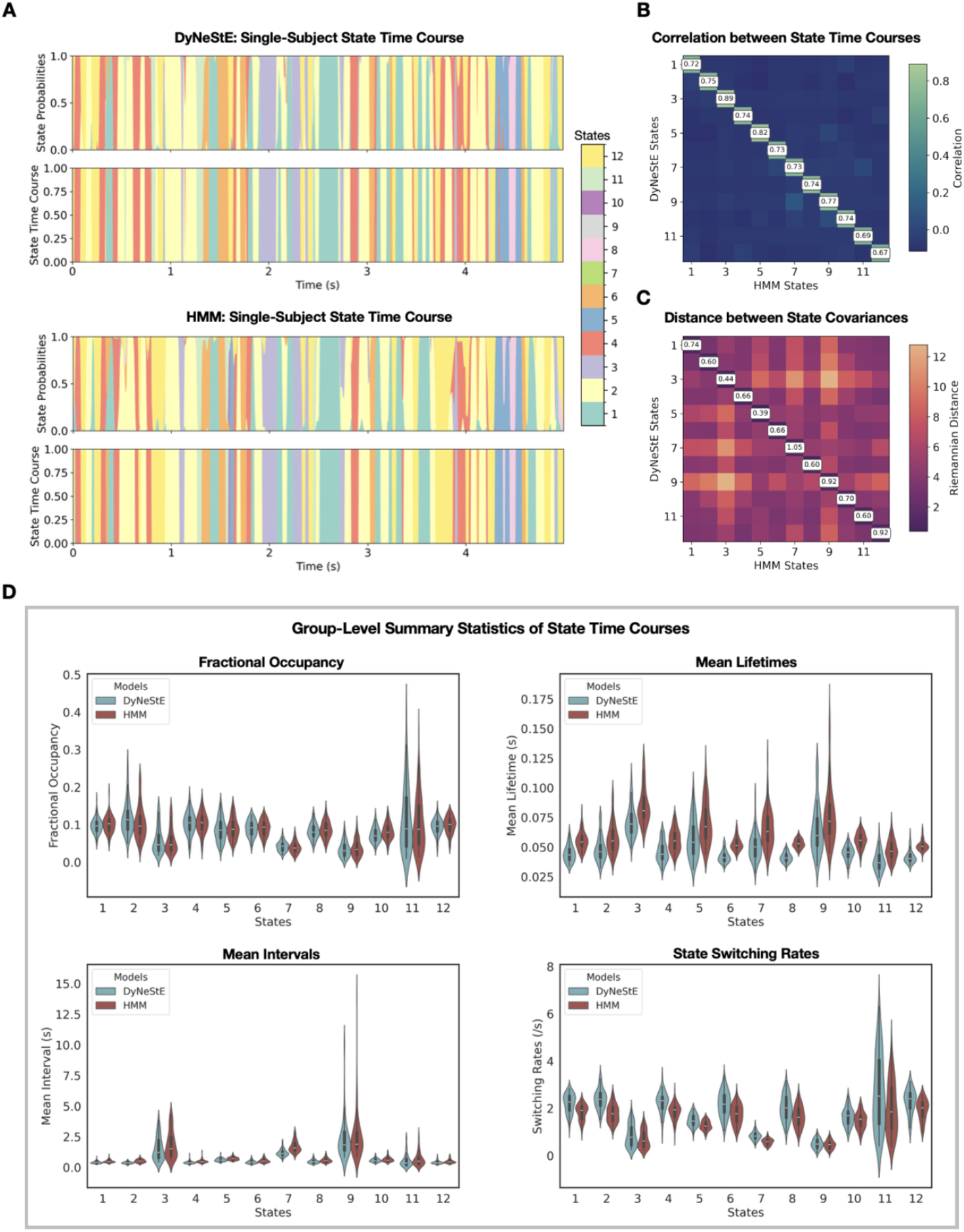
Network dynamics of DyNeStE and HMM state activations. **(A)** State probability traces (top) and state time courses (bottom) inferred by DyNeStE and the HMM for a single subject, shown over the first 5 seconds. **(B)** Pairwise Pearson correlations between inferred state time courses of DyNeStE and the HMM, averaged across subjects. **(C)** Pairwise Riemannian distances between group-level covariance matrices inferred by DyNeStE and the HMM. **(D)** Four group-level summary statistics of DyNeStE (blue) and the HMM (red), depicted as violin plots with box plots overlaid. In all subplots, state ordering follows the networks presented in Figure 3.

To understand potential differences in the state transitions learnt by each model, we subsequently examined summary statistics computed from the respective state time courses (Fig. 4D). Both DyNeStE and the HMM yielded highly comparable fractional occupancies, indicating similar overall proportions of state activation. However, DyNeStE states exhibited shorter mean lifetimes, shorter mean intervals, and higher switching rates across all states. These findings suggest that DyNeStE captures more transient states that activate more frequently compared to the HMM. Importantly, DyNeStE identified a long mean interval of the posterior default mode network (State 9), which is a key feature previously associated with HMM-based modelling of the resting-state network dynamics (Baker et al., 2014).

In summary, DyNeStE successfully inferred canonical, categorical dynamic RSNs comparable to those obtained with the HMM. Both models produced highly similar network profiles and discrete state transitions. However, the state dynamics were not identical, with DyNeStE showing slightly higher switching rates.

#### 3.2.2 Brain networks inferred by DyNeStE are reproducible across data subsets

Because DyNeStE is a stochastic model with a non-convex objective function, it is susceptible to local minima and, consequently, to high variability in its inference. To ascertain whether the RSNs inferred by DyNeStE are reproducible, we divided the dataset into two subsets of subjects and assessed whether the model recovers consistent network descriptions from each half. We found that all states derived from the split datasets were comparable to the RSNs inferred from the full dataset, except for State 7 (Fig. 5). Although minor differences were observed between the two halves, a qualitative comparison of their power maps revealed comparable spatial patterns of power values. To quantify reproducibility, we also computed cosine similarities between corresponding state power maps and tested their statistical significance using a max-statistic permutation test. All states, except State 7, exhibited statistically significant reproducibility (Fig. 5, bottom).

**Figure 5:**
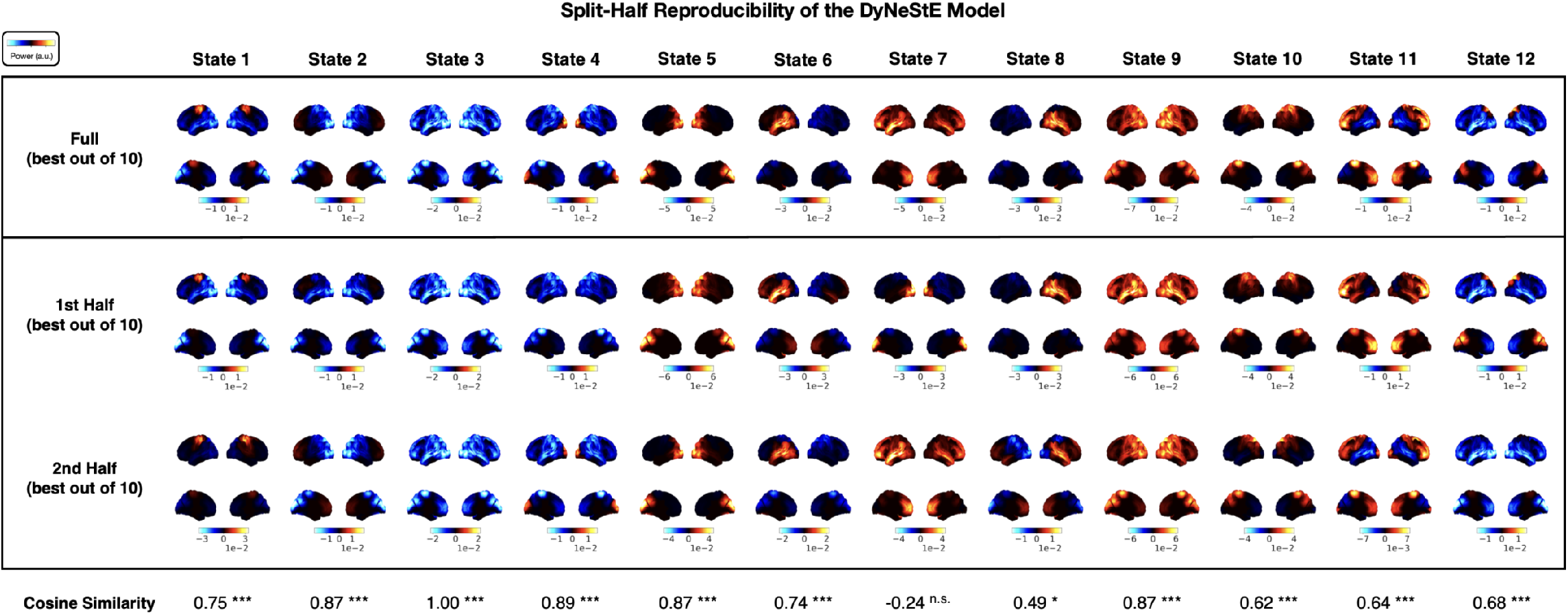
Reproducibility of DyNeStE resting-state networks across split-halves. DyNeStE was trained on two split-halves of the Nottingham MEGUK dataset, with the best model (out of 10 runs) selected for each subset. Group-level power maps inferred from the full dataset (top), the first split-half (middle), and the second split-half (bottom) are shown, each displayed relative to the mean power across all states. At the bottom, cosine similarities between split-half power maps are reported for each state, with statistical significance assessed using a max-statistic permutation test over the states (***: *p <* 0.001, **: *p <* 0.01, *: *p <* 0.05, n.s.: non-significant).

The reduced similarity observed for certain states may arise from non-identifiability, i.e., situations where multiple solutions fit the data equally well. This issue can be exacerbated when the dataset size is limited, thereby increasing the influence of inter-subject variability. To investigate this possibility, we repeated the analysis by training DyNeStE with varying numbers of states (Fig. A6). When the model was trained with 4, 6, or 8 states, all inferred networks were statistically reproducible across split halves. On the other hand, as with 12 states, reproducibility was not consistently observed when training with 10 states. These results suggest that RSNs inferred by DyNeStE are reproducible when the number of states is low enough, given a particular dataset. Similarly, larger datasets are expected to provide reproducibility in models with a greater number of states.

The constraint given by the number of states, however, is expected to diminish with larger datasets, where increased sample size reduces variability and thereby improves statistical power.

#### 3.2.3 DyNeStE learns long-range temporal dependencies

After confirming that DyNeStE recovers canonical and reproducible dynamic RSNs comparable to those identified by the HMM, we next asked whether DyNeStE captures any long-range temporal dependencies via its model parameters that describe the temporal dynamics (i.e., the inferred model RNN). To investigate this capacity, we employed two complementary metrics: the Fano factor and time-lagged mutual information (MI).

As described in Section 2.9.1, the Fano factor measures whether state activations are regularly distributed over time or temporally clustered. For the inferred state time courses of DyNeStE and the HMM, Fano factors started at the value near 1 and increased as a function of window length (Fig. 6A, left). This growth indicates that state activations became increasingly irregular at longer timescales, hinting at the presence of long-range temporal dependencies rather than regular, memoryless state activities. Here, DyNeStE exhibited a slightly steeper increase in Fano factors, with a max-t cluster permutation test over time windows reporting significant differences between the two models at window lengths of approximately 0.1-1 s.

**Figure 6:**
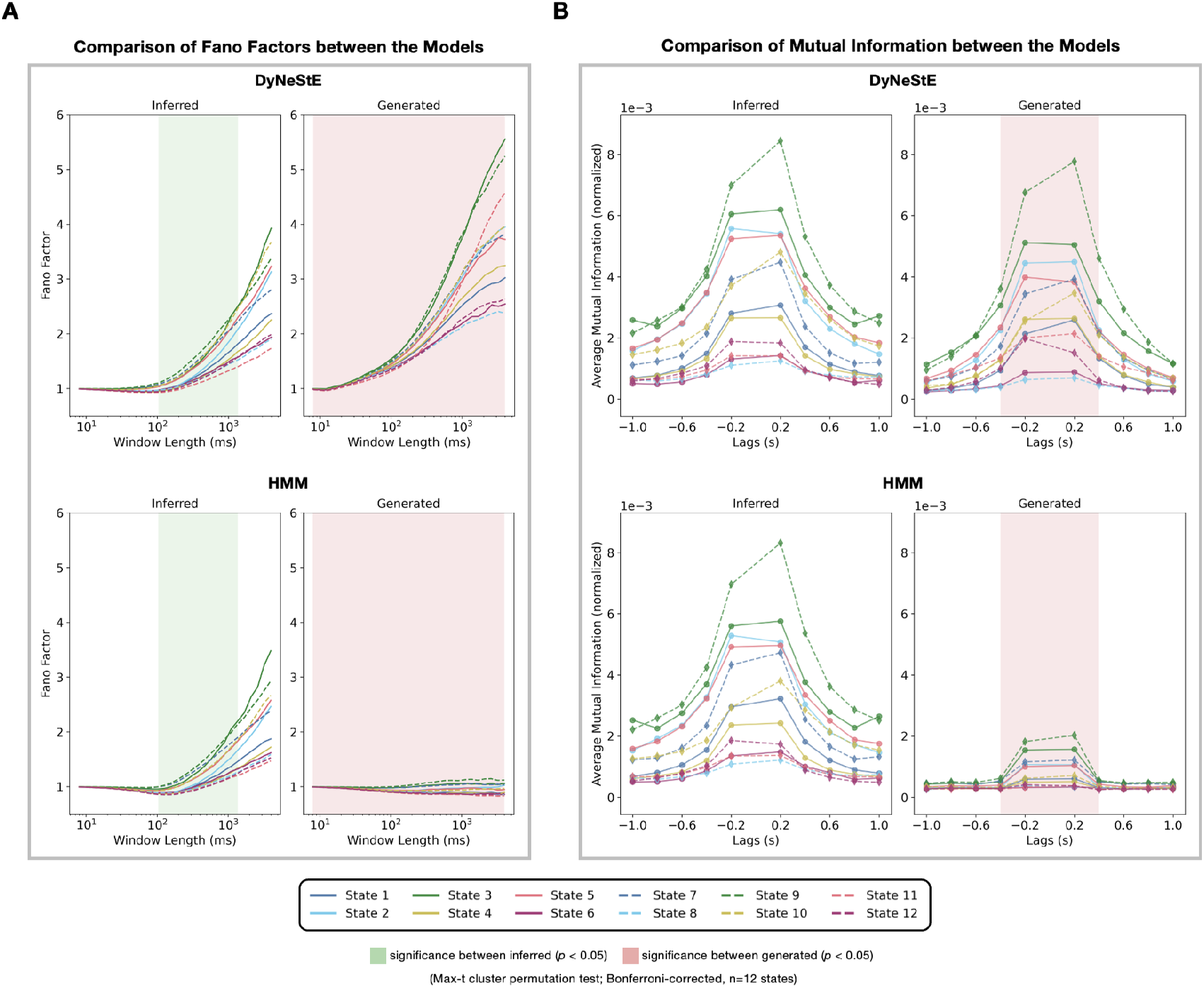
DyNeStE captures long-range temporal dependencies in real MEG data. DyNeStE’s ability to represent long-term dependencies was quantified using Fano factors and mutual information. **(A)** Subject-averaged Fano factors, computed from both inferred and generated state time courses, are shown for DyNeStE (top) and the HMM (bottom). Each line corresponds to Fano factors of each state across window lengths. Time windows showing significant group-level differences between the two models (*p <* 0.05; max-t cluster permutation test over time dimension, Bonferroni-corrected, *n* = 12 states) for all states are marked by green (inferred) and red (generated) shadings. **(B)** Normalised mutual information between each state and all other states, computed from both inferred and generated state time courses, are shown for DyNeStE (top) and the HMM (bottom) across temporal lags. Visualisation and statistical testing procedures are identical to those described in (A).

However, patterns observed in the inferred state time courses may purely reflect data-driven information, with the generative models not capturing any long-range dependency information in their modelling of the temporal dynamics (i.e., the transition probability matrix in the HMM and the model RNN in DyNeStE). To investigate this, rather than using the inferred state time courses, we generated state time courses from the inferred DyNeStE and HMM generative models and recomputed Fano factors (Fig. 6A, right). Consistent with our simulation result (Fig. 2C), DyNeStE preserved the characteristic increase in Fano factors with longer window lengths, whereas the HMM produced values near 1 across all window lengths, showing that long-range temporal dependencies have been captured by DyNeStE but not the HMM. Across the full range of window lengths, a max-t cluster permutation test revealed significant differences between the two models.

To consolidate this finding, we also examined the time-lagged MI, which quantifies statistical dependencies between network states, specifically the extent to which a state carries information about past or future activations. For both DyNeStE and the HMM, the time-lagged MI calculated using the inferred state time courses exhibited higher values at short lags that decayed gradually as the lag increased (Fig. 6B, left), suggesting that their state activations retain predictive information across extended timescales. Unlike the Fano factors, however, no significant differences between the models were observed. In contrast, when the time-lagged MI was computed using the generated state time courses (Fig. 6B, right), HMM-derived states showed greatly reduced MI at short lags and near-zero MI at longer lags, whereas DyNeStE maintained MI values comparable to those computed from its inferred state time courses. These differences were significant at temporal lags of approximately −400 to +400 ms. Although not statistically significant, DyNeStE also exhibited a more gradual MI decay at longer lags compared to the HMM.

Taken together, the analyses of Fano factors and time-lagged MI from both observed and generated data demonstrate that DyNeStE is able to capture long-range temporal dependencies, whereas the HMM does not. This finding underscores an advantage over the HMM, as DyNeStE retains a longer memory of state histories, which can provide a stronger temporal prior when inferring states.

#### 3.2.4 DyNeStE captures long-term cyclic structures of the dynamic brain networks

A previous study using the HMM has shown that sequential activations of brain networks are organised in cycles during rest, particularly when state activations are separated by long intervals (van Es, Higgins, et al., 2025). Here, we tested whether DyNeStE could similarly infer such long-term cyclical pattern and represent it through its learnt model parameters.

We first applied TINDA (see Section 2.10) to the inferred state time courses of DyNeStE and the HMM trained on the Nottingham MEGUK dataset. Consistent with the findings of van Es, Higgins, et al. (2025), both models revealed cycles with identical state ordering from their state transitions (Fig. 7A). However, when TINDA was applied to the state time courses *generated* from the learnt models, rather than to the inferred state time courses, the cyclical structure disappeared for the HMM but persisted for DyNeStE (Fig. 7B). This implies that only DyNeStE can incorporate the long-term cyclical organisation of state transitions in its representations. This discrepancy was also quantitatively confirmed, with DyNeStE showing stronger cycle strength than the HMM (Fig. 8A).

**Figure 7:**
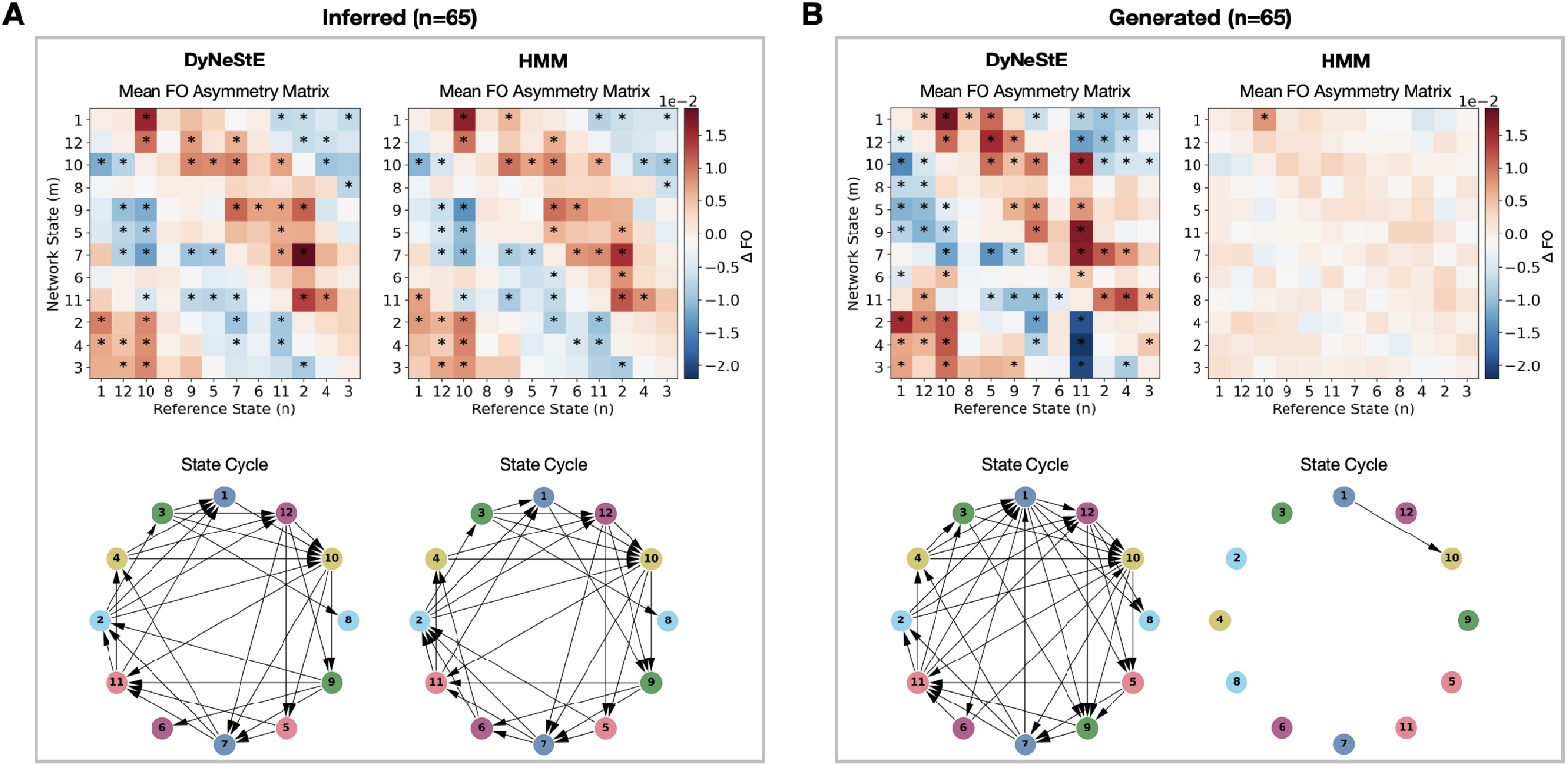
Cyclical organisation of state activations in observed and generated data [van Es, Higgins, et al. (2025)]. **(A)** FO asymmetry matrices (top), computed from the inferred state time courses of DyNeStE and the HMM, averaged across 65 Nottingham MEGUK participants. These matrices summarise whether a given reference state is more likely to precede or follow other network states. The state cycles derived from these matrices are shown below. Statistically significant edges, identified using a paired-samples t-test (*p <* 0.05, Bonferroni-corrected with *n* = 132 state pairs; see Section 2.10.3), are marked with asterisks in the group-averaged FO asymmetry matrices and visualised as arrows in the state cycles. **(B)** Identical to (A), but using the generated state time courses.

**Figure 8:**
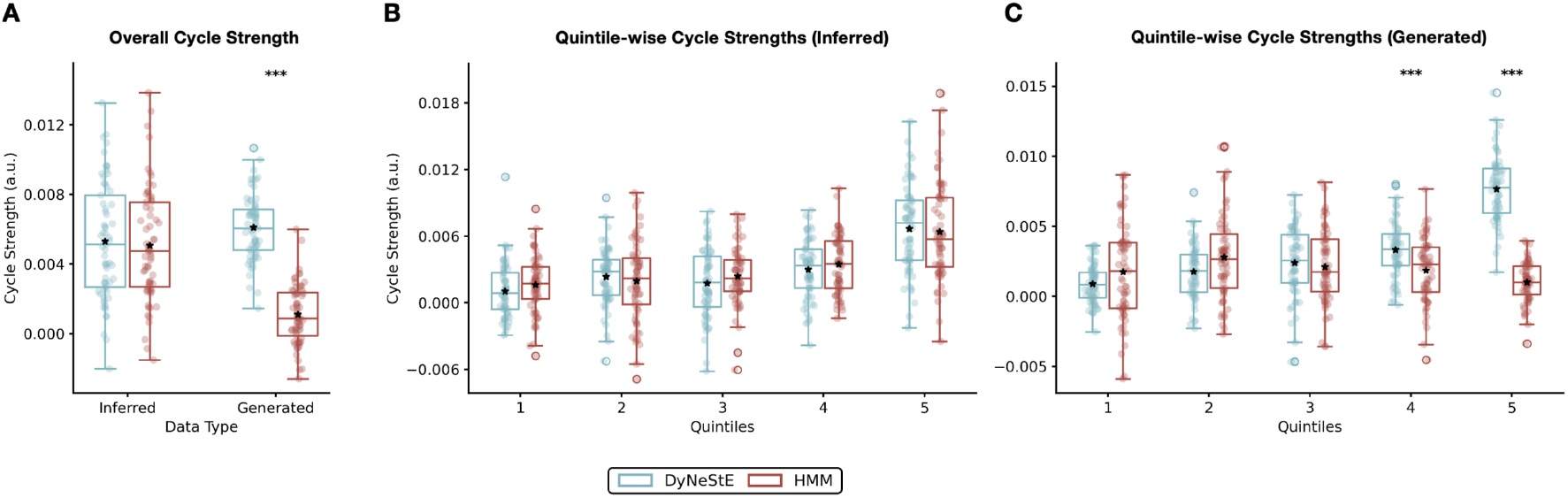
Cycle strengths of state transitions in observed and generated data. **(A)** Overall cycle strengths, quantifying the presence of cyclical structure underlying dynamic brain networks, were computed from inferred and generated state time courses of DyNeStE (blue) and HMM (red) trained on the Nottingham MEGUK dataset. Box plots show the distribution across subjects, with black stars indicating the mean. Group differences were assessed using a two-samples t-test. **(B)** Quintile-wise cycle strengths were computed by dividing inter-state intervals from inferred state time courses into five bins and estimating cycle structure separately within each bin. Box plots show the distribution across subjects, with means indicated by black stars. Group differences were tested using a Welch’s t-test with Bonferroni correction (n=5 quintiles). **(C)** Same as (B), but using generated state time courses. Statistical significance is indicated by asterisks (***: *p <* 0.001, **: *p <* 0.01, *: *p <* 0.05); non-significant results are not marked.

Because the previous work reported that such cyclical organisation is primarily driven by long ISIs (c.f., Figure 3 in van Es, Higgins, et al. (2025)), we also stratified the generated state time courses into five quintiles based on the ISIs (Fig. 9A) and repeated the TINDA analysis within each bin. In line with the results above, while DyNeStE successfully recovered a cyclical structure from the generated data, the HMM did not^15^ (Fig. 9B-D). As expected, this cyclical organisation was only evident in the fifth quintile, corresponding to the longest ISIs. For inferred state time courses, both models revealed a clear cycle in the fifth quintile (Fig. A7). This observation was supported by the statistical tests, with DyNeStE reporting significantly stronger cycle strengths compared to the HMM for the generated data (Fig. 8B-C).

**Figure 9:**
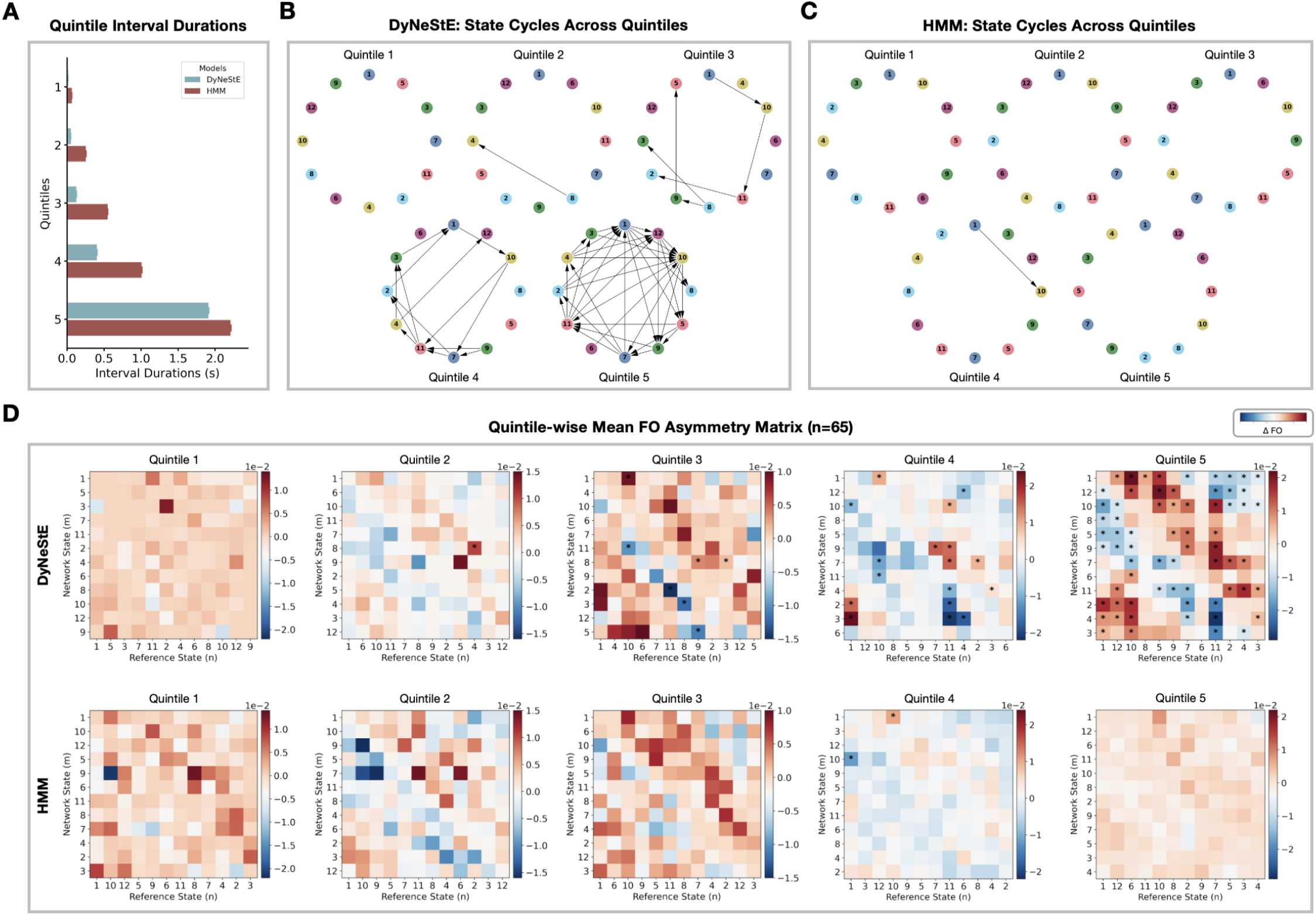
Quintile-wise cyclical organisation of state activations in generated data. **(A)** Distribution of inter-state interval durations across quintiles, with mean values shown as bars and standard errors as error bars, for DyNeStE (blue) and the HMM (red). **(B)** Quintile-specific state cycles derived from the state time courses generated by DyNeStE. **(C)** Quintile-specific state cycles derived from the state time courses generated by the HMM. **(D)** Group-averaged FO asymmetry matrices for each quintile, shown for DyNeStE (top) and the HMM (bottom). Asterisks mark statistically significant edges identified using a paired-samples t-test (*p <* 0.05, Bonferroni-corrected with *n* = 132 state pairs).

In addition, it is noteworthy that in Figure 7, the order of States 5 and 9 is swapped between the inferred and generated cycles for DyNeStE. However, a closer inspection of the fifth-quintile cycles (Fig. 9B, Fig. A7B) shows a consistent ordering across these cycles, suggesting that the discrepancy is likely attributable to noise introduced by state transitions associated with shorter ISIs.

In summary, while both DyNeStE and the HMM replicated the cyclical organisation of dynamic brain networks in inferred state transitions, only DyNeStE preserved this structure in its generative data. This finding reinforces our conclusion in Section 3.2.3, as it demonstrates that DyNeStE not only captures long-range temporal dependencies during inference but also embeds them within its generative process by encoding these dependencies in its model parameters.

### 3.3 Real Data Analysis: Replay

Lastly, having evaluated DyNeStE’s performance on the Nottingham MEGUK dataset, we wanted to confirm whether the results above are reproducible on a new, independent dataset. To this end, we applied DyNeStE to the Replay dataset used in Higgins et al. (2021). Because the dataset does not include sMRI scans of the participants^16^, we utilised the pre-trained DyNeStE and HMM models, applying them directly to infer state time courses without further training (i.e., with model parameters fixed to those learnt from the Nottingham MEGUK dataset).

Based on these state time courses, we tested if DyNeStE can learn categorical states of neural dynamics and the underlying long-range temporal dependencies sufficiently well to replicate prior findings. Looking at the spatio-spectral descriptions of the RSNs (Figs. A8, A9), we first verified that the inferred networks are comparable between DyNeStE and the HMM, as observed in the Nottingham MEGUK dataset. Nonetheless, due to the lack of sMRI scans, their FC networks appeared somewhat noisier.

In Higgins et al. (2021), the authors used the HMM to demonstrate that memory-associated replay bursts and RSNs share a common long-range temporal dependency. Consistent with this result, we observed that both DyNeStE and the HMM capture this structure, as evidenced by replay-evoked network activations (Fig. 10A-B, left). Specifically, States 5, 7, and 9 (visual, anterior default mode, and posterior default mode networks) showed concordant activations with replay bursts, replicating the previous findings (c.f., Figure 2 in Higgins et al. (2021)). State-wise replay intervals and rates (Fig. 10A-B, middle) further indicated that these states were associated with longer replay intervals and higher replay rates, although State 7 in DyNeStE showed statistically insignificant replay-interval specificity (c.f., Figure 3 in Higgins et al. (2021)).

**Figure 10:**
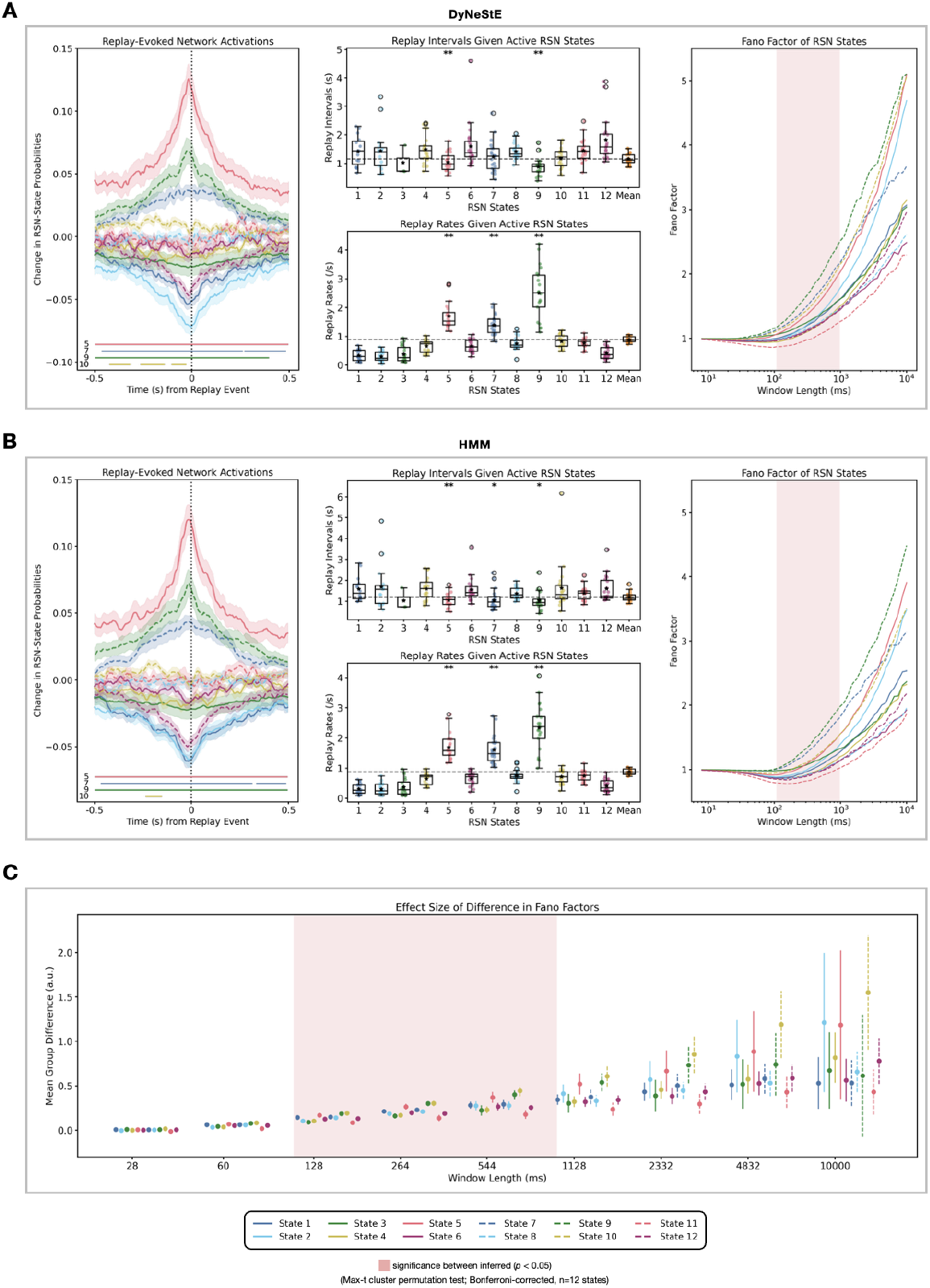
DyNeStE reveals long-term temporal structure shared between replay bursts and RSNs. **(A)** Replay-evoked network activations (left) show changes in state probabilities around replay onsets. Statistically significant time points are indicated by coloured lines labelled with the corresponding state number, following a max-t cluster permutation test over the time dimension(*p <* 0.05). Replay intervals and rates associated with each RSN state (middle) are shown, with the overall mean across states depicted as a gray dotted line. For each state, significance was assessed with a one-sample max-t permutation test across subjects on the respective replay metric (***: *p <* 0.001, **: *p <* 0.01, *: *p <* 0.05). Fano factors (right) were computed from DyNeStE-inferred state time courses. Red shading marks window lengths where DyneStE significantly differed from the HMM for all states, following a max-t cluster permutation test over time windows (*p <* 0.05). All p-values were Bonferroni-corrected for the number of states (n=12). **(B)** Same analyses as in (A), but applied to state time courses inferred by the HMM on the Replay dataset. **(C)** Effect sizes of the differences in Fano factors between DyNeStE and the HMM, computed as the mean group-level difference. Error bars denote the standard error of the mean group difference between the models, and red shading indicates window lengths with significant differences between the two models across all states (*p <* 0.05; Bonferroni-corrected, n=12 states).

Finally, the Fano factors of the state time courses inferred by both DyNeStE and the HMM indicated the presence of long-range temporal dependencies in state transitions (Fig. 10A-B, right). Notably, in line with Section 3.2.3, DyNeStE exhibited larger Fano factors than the HMM (Fig. 10C), exhibiting a stronger expression of these dependencies.

In summary, DyNeStE reliably inferred categorical states and replicated HMM-based findings on replay-related network dynamics. Moreover, DyNeStE detected stronger long-range temporal dependencies from the network state transitions than the HMM, as evidenced by its higher Fano factors.

## 4 Discussion

In this paper, we proposed DyNeStE as a new methodology for modelling dynamic brain networks. The contributions of this model are twofold. First, it represents dynamic brain networks as categorical latent states. Using both simulated and real MEG data, we have shown that DyNeStE can infer plausible RSNs (Figs. 2A-B, 3, A8), which are comparable to those of the HMM (Figs. 4, A4, A9) and reproducible across independent subsets of the Nottingham MEGUK dataset (Figs. 5, A6). Secondly, in both simulated and real MEG datasets, DyNeStE successfully captured long-range temporal dependencies in its learnt model parameters, whereas the HMM could only infer such effects in a data-driven manner (Figs. 2C, 6-10). Thus, DyNeStE substantially enhances the ability to investigate the activity of large-scale brain networks at long timescales.

### 4.1 Advantages of DyNeStE

In DyNeStE, brain network dynamics are modelled as discrete state transitions by constraining the posterior distribution to be categorical. Additionally, to make the model capture long-range temporal dependencies in state dynamics, we dropped the assumption that network dynamics follow a first-order Markovian process by implementing a recurrent neural network (i.e., LSTM). These attributes position DyNeStE as a strong alternative to current state-of-the-art approaches. While HMMs are restricted to modelling Markovian dynamics, existing neural network-based approaches such as DyNeMo, which encode states in a continuous latent space, allow multiple states to co-occur at the same time point, potentially compromising interpretability (Gohil et al., 2022). Conversely, DyNeStE can generalise the HMM by converging to HMM-like solutions when the underlying dynamics are categorical and extend its capabilities to incorporate rich temporal structures when long-range temporal dependencies are present in the data. In this way, DyNeStE addresses the existing limitations, potentially yielding a more adaptive characterisation of the data.

It can be asserted that the HSMM offers a potential alternative for modelling long-term dependencies, as it appears to address a similar limitation as DyNeStE (Trujillo-Barreto et al., 2024). However, HSMMs typically require the distribution of state duration to be specified explicitly, and even if this requirement can be relaxed through nonparametric formulations, the model remains bound by the first-order Markovian assumption. Hence, dependencies are confined to the immediately preceding state: memory is extended only through state persistence (i.e., duration), rather than by capturing long-term dependencies across distinct states. To this extent, HSMMs cannot leverage past state history to model complex temporal dynamics, leaving them inherently restricted compared to the recurrent approaches (i.e., RNNs) employed in DyNeStE.

By adopting categorical representations, however, DyNeStE is implicitly assuming that brain networks are mutually exclusive. One may argue that this assumption is overly restrictive. For instance, previous work has shown that modelling such co-activation patterns by relaxing the mutual exclusivity constraint with DyNeMo explains task data better (Gohil, Kohl, Huang, et al., 2024). Nevertheless, improvements in describing spatio-spectral dynamics through this relaxation were reported to be modest in DyNeMo relative to the HMM, as evidenced by marginal differences in reconstruction error for task-dependent spectrograms between the two models (Gohil et al., 2022). Given the comparability between DyNeStE and the HMM in their inferred dynamics (Figs. 3-4), we expect the same conclusion to hold. Thus, while network co-occurrence is possible, DyNeStE opts for a trade-off, choosing interpretability and robustness over modelling simultaneous network activations.

### 4.2 Comparative Validation of DyNeStE and HMM

Another important validation outcome in our study is that DyNeStE successfully replicates key findings from two HMM-based studies (Higgins et al., 2021; van Es, Higgins, et al., 2025). This replication demonstrated that DyNeStE can preserve the strengths of the HMM framework while extending its capabilities, being able to learn long-range temporal dependencies robustly across multiple datasets (Figs. 6-10). Viewed from another perspective, our findings can also be interpreted as reinforcing the validity of the HMM itself. Apart from the advantage in modelling long-term dynamics noted above, DyNeStE and the HMM produced generally consistent results across all the analyses performed on inferred network states. Their similarity highlights the robustness of data-driven inference^17^ by the HMM despite its short-term memory.

### 4.3 Applications and Generative Modelling

The ability of DyNeStE to explicitly learn long-range dependencies has important implications for both neuroscience and computational modelling. First, it opens opportunities for *interpretability* studies. Because the latent representations of DyNeStE encode temporal structure across longer time spans, they can be used to study how network states interact. This makes DyNeStE promising for task paradigms in which long-term processes such as attention and memory are in question. For example, estimated posterior distributions or state observation models could be examined to identify networks supporting specific cognitive abilities. One practical approach would be to derive subject-specific covariance matrices from group-level estimates using *dual estimation* (Vidaurre et al., 2021) and assessing whether these covariances can predict the long-term cognitive performance of each subject.

Another prospective application of DyNeStE lies in its data generative capacity. As a generative model for electrophysiological data, its architecture naturally supports the simulation of signals that incorporate long-term dependencies and cyclical structure, enabling it to produce more realistic MEG-like activity than models lacking such capacity. This usability could be valuable for simulation studies aiming to probe how local cellular mechanisms relate to macroscopic brain activity or how animal electrophysiology connect to human neuroimaging data.

This generative application also extends to the field of machine learning (ML) and artificial intelligence. Deep learning studies using neuroimaging data often suffer from data scarcity, compounded by privacy issues, especially regarding the sharing of clinical recordings (Darvishi-Bayazi et al., 2024; Kinahan et al., 2024; Yeom et al., 2023; Zhu et al., 2024). Given the limited availability of open M/EEG datasets, synthetic data generation has been proposed as a promising method, both for data augmentation and privacy-preserving data releases^18^ (Lu et al., 2025; Torma & Szegletes, 2025). Furthermore, in a speech decoding task, the scaling law for neural networks has been shown to hold with MEG data (Jayalath et al., 2025). With larger dataset, generalisation across participants, datasets, and tasks became possible, and the model performance matched the one obtained using invasive electrophysiological recordings. Under these circumstances, DyNeStE is well-suited for this application, as it can generate biologically plausible signals that capture realistic network transitions.

### 4.4 Methodological Contributions

Beyond its applications, DyNeStE makes several methodological contributions to VAE architectures. Categorical VAEs with a Gumbel-Softmax distribution have traditionally been limited to evaluations on small-scale datasets and have struggled to match the performance of continuous VAEs that rely on the Gaussian reparametrisation trick (van den Oord et al., 2017). To our knowledge, DyNeStE is one of the first successful applications of a categorical VAE with Gumbel-Softmax to large-scale, real neural datasets, substantially narrowing this performance gap. In particular, DyNeStE trains as robustly as DyNeMo, which serves as a continuous latent counterpart. Moreover, it employs a learnt categorical prior (rather than a uniform fixed prior), enhancing its flexibility and expressiveness. These characteristics highlight the potential of DyNeStE to be utilised as a foundation model that provides regularisation for smaller-scale models (e.g., through knowledge distillation or transfer learning), in which limited data availability makes training categorical latent variable models from scratch especially challenging.

### 4.5 Methodological Limitations

Despite its strengths, DyNeStE presents several methodological limitations pertaining to its architectural design and optimisation.

#### 4.5.1 Transient State Activations

The state time courses inferred and generated by DyNeStE typically include very fast state activations. As shown in Figure 4, DyNeStE infers states with lifetimes shorter than those detected by the HMM. Two non-exclusive factors may contribute to this transiency: (i) stochasticity introduced by the Gumbel-Softmax sampling during training and (ii) DyNeStE’s enhanced sensitivity to transient neurophysiological changes afforded by its long-range temporal modelling. If the former dominates, the transient state activations may reflect noise in the data. While modelling uncertainty is inherent to a variational framework and considered beneficial, excessive transient states could reduce the accuracy of generated data and complicate the interpretation of state time courses in this case.

#### 4.5.2 Computational Demands and Training Complexity

DyNeStE is computationally demanding and more difficult to train than alternative models (i.e., DyNeMo and the HMM). As discussed in Appendix A.3.1, the model optimisation requires careful scheduling of multiple parameters (i.e., KL divergence, Gumbel-Softmax temperature, and learning rate), and the categorical latent space often forms a loss landscape that is non-convex, noisy, and non-smooth. For example, DyNeStE required approximately 350 training epochs^19^ on the Nottingham MEGUK dataset, compared to only 20 for the HMM and 60 for DyNeMo. Thus, while DyNeStE offers richer modelling capacity, its added complexity and computational cost may not be justified in contexts where HMMs provide sufficient inference (see Section 4.2) and long-term dependencies are not of primary interest.

#### 4.5.3 Limits of Recurrent Neural Networks

Although DyNeStE overcomes the short-term memory of HMMs, its recurrent architecture is itself bounded by GPU memory and the vanishing gradient problem, albeit with its use of LSTMs and layer normalisation. Consequently, there are practical limits to the RNN receptive field that can be exploited during training. This was hinted at in the TINDA analysis, where, for the fifth quintile in which ISIs exceeded the RNN sequence length, the inferred cycle structure appeared noisier (Fig. A7). Nevertheless, DyNeStE still captured stronger cycle strength than the HMM, underscoring its relative advantage even under such constraints.

Another limitation stemming from the LSTM architecture is the trade-off between interpretability of learnt temporal structure and the ability to model long-range temporal dependencies. While LSTMs offer a powerful framework for capturing long-term dynamics, their nonlinear and high-dimensional parametrisation hinder direct interpretability of the underlying state transition dynamics. This contrasts with the HMM, which produces an explicit transition matrix outlining state-to-state probabilities. In this paper, we addressed this limitation by evaluating DyNeStE’s ability to capture long-range temporal dependencies through analysing the statistical properties of the synthetic data it generates. Future studies on the mechanistic interpretability of recurrent neural networks may enable the theoretical derivation of principled dynamical descriptions analogous to HMM transition matrices.

#### 4.5.4 State Identifiability

DyNeStE may still face challenges with identifiability in certain configurations. While the mutual exclusivity enforced by DyNeStE can attenuate this issue to some extent, different sets of networks may still explain the data equally well, particularly when using a number of states. For example, in Section 3.2.2, analyses of the 12-state DyNeStE revealed reduced split-half reproducibility for State 7, suggesting that there may be an alternative solution for the dataset. Notably, this same state was not associated with shorter replay intervals, contrary to what was reported in the prior study (Fig. 10A). Reduced split-half reproducibility, however, typically improves with larger datasets and fewer states. This improvement hints that increasing the sample size or reducing the number of states would likely mitigate the identifiability issue.

### 4.6 Outlooks and Future Research Directions

DyNeStE offers a range of opportunities for future research and methodological development. First, the generalisability of DyNeStE should be evaluated across modalities such as EEG and fMRI, as well as across different scanner types and on sensor space data. Demonstrating robustness in diverse data contexts would corroborate confidence in its applicability beyond MEG.

Secondly, applying DyNeStE to task-based and clinical datasets is a promising direction. These datasets may feature network activations that persist over a longer period of time or manifest more structured temporal dependencies, where DyNeStE’s ability to model long-term dynamics may offer benefits. This is supported by a modest but consistent advantage of DyNeStE in capturing long-range temporal dependencies, as reflected by larger effect sizes in the Fano factor analysis (Figs. 6A, 10C). Confirming these advantages in applied settings would underscore the model’s translational relevance.

Thirdly, the generative capacity of DyNeStE can be explored in greater detail. Synthetic data produced by the model could be used to study long-term network dynamics, test theoretical hypotheses on network couplings, or support downstream applications such as data augmentation for machine learning. Assessing the fidelity and utility of these synthetic datasets remains an important avenue of investigation.

Finally, methodological refinements may help reduce training complexity and computational cost. For example, researchers have proposed replacing the categorical prior with a Gumbel–Softmax prior, or applying exponential transformations to the Gumbel–Softmax distribution to represent discrete random variables in log space, as potential strategies to prevent gradient underflow and stabilise optimisation (Maddison et al., 2017). Systematic testing of such strategies could make DyNeStE more accessible for broader use.

## 5 Conclusion

We introduce DyNeStE, a novel generative model that represents brain networks as categorical states and captures long-term dynamics in MEG data using an RNN architecture. Our findings demonstrate that DyNeStE scales efficiently to large MEG datasets, replicates established HMM-based findings, and trains robustly across multiple datasets. Overall, DyNeStE offers an alternative to current methods, with the potential benefits of more accurate modelling and generation of brain network dynamics.

## Data and Code Availability

The Nottingham MEGUK dataset is publicly available at https://meguk.ac.uk/database/ (raw sensorlevel MEG recordings) and at https://osf.io/by2tc/ (source reconstructed data). The Replay dataset is freely available upon request, subject to participant consent (Higgins et al., 2021; Liu et al., 2019).

The source code for DyNeStE can be found in the osl-dynamics toolbox (Gohil, Huang, et al., 2024), along with example scripts for basic usage. All scripts for reproducing the results, figures, and analyses in this paper are written in Python and available at https://github.com/OHBA-analysis/Cho2026_DyNeStE.

## Ethics Statement

The Nottingham MEGUK dataset was collected at the University of Nottingham, UK, as part of the UK MEG Partnership. All participants gave written informed consent, and ethical approval was obtained from the University of Nottingham Medical School Research Ethics Committee. The replay dataset was collected at the University College London (UCL), UK, with participants recruited from the UCL Institute of Cognitive Neuroscience subject pool. All participants provided written informed consent prior to participation, and the study was approved by the UCL Research Ethics Committee (ethics number 9929/002).

## Use of Generative Artificial Intelligence Assistance

In preparing this manuscript, the authors used artificial intelligence (AI) tools to assist with language editing, grammar correction, and stylistic refinement of select sections. All changes were reviewed, verified, and edited by the authors. No new content was generated by AI tools, and the authors take full responsibility for the content of the publication.

## Author Contributions

S.C.: Conceptualisation; Methodology; Software; Validation; Formal Analysis; Data Curation; Writing – Original Draft; Writing – Reviewing & Editing; and Visualisation. R.H.: Conceptualisation; Methodology; Software; Writing – Reviewing & Editing. C.G.: Conceptualisation; Methodology; Software; Data Curation; Writing – Reviewing & Editing. O.P.J.: Methodology; Writing – Reviewing & Editing; Supervision. M.W.W.: Conceptualisation; Methodology; Data Curation; Writing – Reviewing & Editing; Supervision.

## Declaration of Competing Interests

The authors declare no competing financial interests or personal relationships that could be perceived as influencing the findings reported in this article.

## Acknowledgements

This research was supported by the National Institute for Health Research (NIHR) Oxford Health Biomedical Research Centre. SC is supported by the Medical Sciences Graduate School Studentship, funded by the Medical Research Council (MR/W006731/1), the Hertford Claire Clifford Lusardi Scholarship, and the Nuffield Department of Clinical Neurosciences. RH is supported by the EPSRC Centre for Doctoral Training in Health Data Science (EP/S02428X/1). CG is supported by the Wellcome Trust (215573/Z/19/Z). OPJ is supported by the Medical Research Council (MR/X00757X/1), Royal Society (RG/R1/241267), National Science Foundation (2314493), NFRF (NFRFT-2022-00241), and SSHRC (895-2023-1022). MW is supported by the Wellcome Trust (106183/Z/14/Z, 215573/Z/19/Z), the New Therapeutics in Alzheimer’s Disease (NTAD) study supported by the Medical Research Council, the Dementia Platform UK (RG94383/RG89702) and the NIHR Oxford Health Biomedical Research Centre (NIHR203316). The views expressed are those of the authors and not necessarily those of the NIHR or the UK Department of Health and Social Care.

## A Appendix

### A.1 Generative Models of DyNeStE and HMM

Here, we compare the generative models of DyNeStE and the HMM. As illustrated in Figure A1, both models share the same observation model, wherein each state is associated with a multivariate normal distribution parametrised by a mean vector and covariance matrix. These models, however, differ in how they model the temporal dynamics. The HMM generates state time courses according to a transition probability matrix learned via the Baum-Welch algorithm with closed-form update rules. In contrast, DyNeStE employs an LSTM network to capture temporal dependencies and learn flexible representations of state transition dynamics, from which state time courses are sampled.

**Figure A1:**
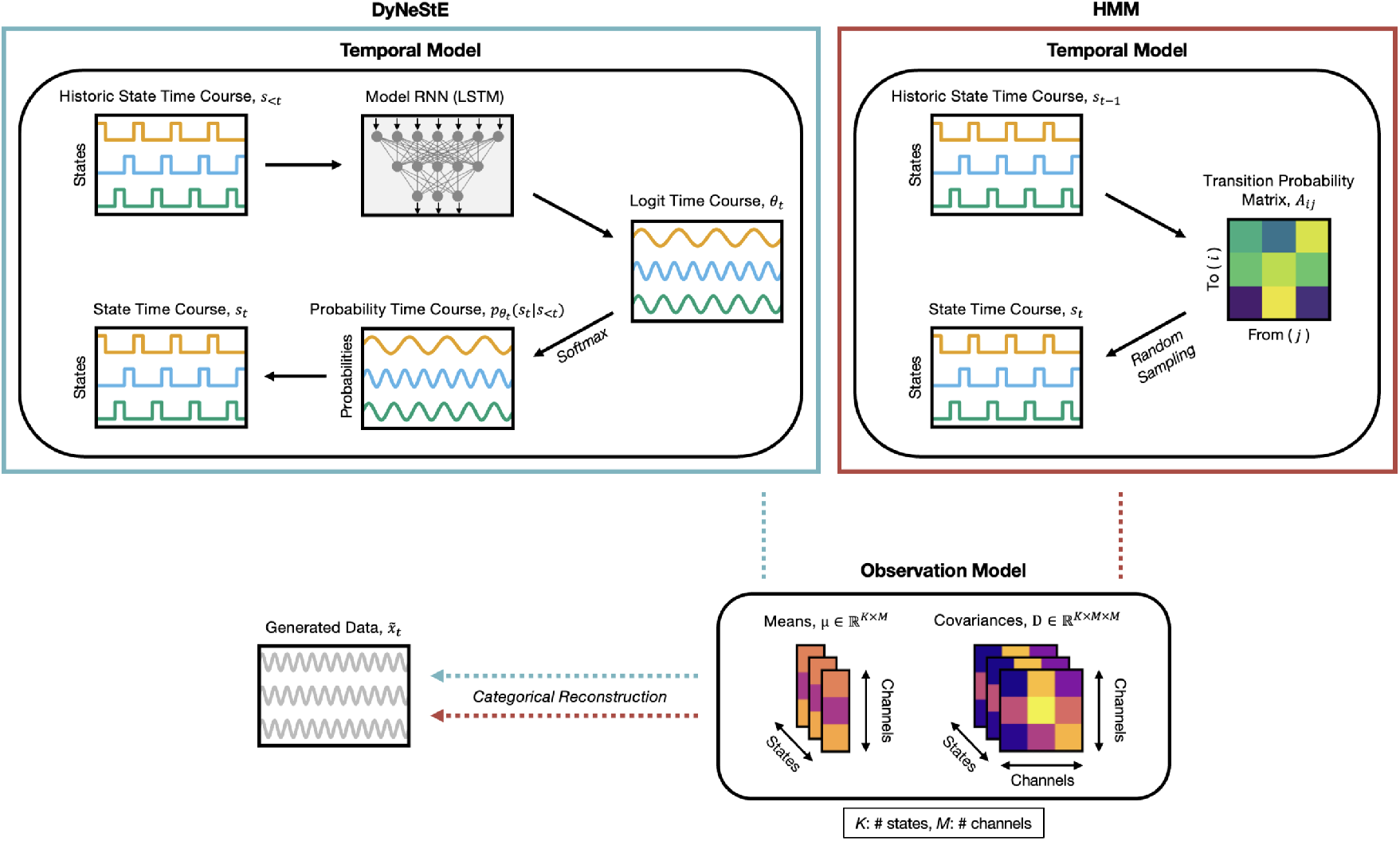
Generative models of DyNeStE and HMM. Generative frameworks for DyNeStE (blue) and the HMM (red) are shown. In DyNeStE, the historic states *s*_*<t*_ are input to an LSTM network, which outputs logits that parametrise a categorical distribution over the current state variable *s*_*t*_. The current state is then sampled from this distribution to generate the state time course. In the HMM, the previous state *s*_*t*−1_ serves as an index to the transition probability matrix, from which the next state *s*_*t*_ is randomly sampled based on the corresponding transition probabilities. Once a state *s*_*t*_ is determined, the observation model (identical for both approaches) for a specific state is selected, and the observed data *x*_*t*_ is generated from the multivariate Gaussian distribution with state-specific means and covariances.

### A.2 Derivation of the DyNeStE Loss Function

In variational Bayesian inference, the objective is to minimise the **variational free energy**, denoted by ℱ:

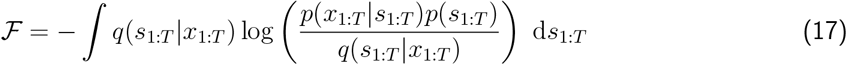

Here, *q*(*s*_1:*T*_ |*x*_1:*T*_ ) is the variational posterior distribution, *p*(*s*_1:*T*_ ) is the prior, and *p*(*x*_1:*T*_ |*s*_1:*T*_ ) is the data likelihood. The observed data *x*_*t*_ is a vector of length *M*, where *M* is the number of channels. The latent state vector *s*_*t*_ is of length *K*, where *K* is the number of hidden states. The index *t* = 1, …, *T* denotes uniformly spaced time points.

The variational free energy can be decomposed into the sum of the negative LL and KL divergence such that

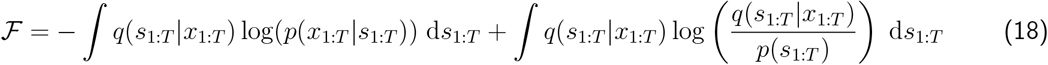

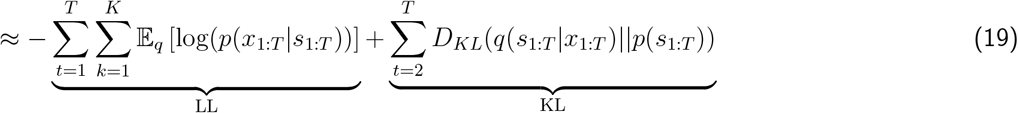

where 𝔼 (·) is the expectation with respect to the posterior *q*(*s*_1:*T*_ |*x*_1:*T*_ ), and *D*_*KL*_(·||·) is the KL divergence.

#### A.2.1 Log Likelihood Loss

Let’s first consider the expected LL loss,

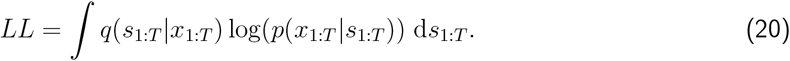

To simplify this term, two mathematical assumptions are employed.

**First**, we apply the mean field approximation to the **posterior** as below:

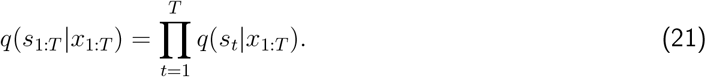

Each *q*(*s*_*t*_|*x*_1:*T*_ ) is modelled as a categorical distribution parametrised by a logit vector:

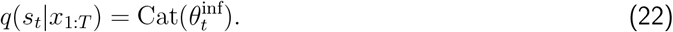

The logits 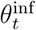 are inferred using a bidirectional LSTM network (i.e., the inference RNN). This network outputs parameters of the variational posterior distribution given the observed data as

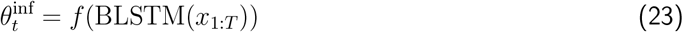

where *f* is a learnable linear affine transformation. Because sampling from the categorical distribution is non-differentiable, we use the continuous Gumbel-Softmax distribution to approximate the categorical distribution by

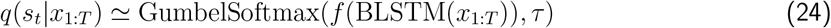

where *τ >* 0 is the temperature of the softmax function. As *τ* → 0, the distribution becomes sharp, and the samples approach one-hot vectors; as *τ* → ∞, the distribution becomes more uniform (c.f., Figure 1 in Jang et al. (2017)).

The posterior *q*(*s*_*t*_|*x*_1:*T*_ ) is now a probabilistic approximation of a one-hot vector, containing the probability of each state. Hence, each element in *q* = (*q*_1_, …, *q*_*k*_) is a Gumbel-Softmax sample

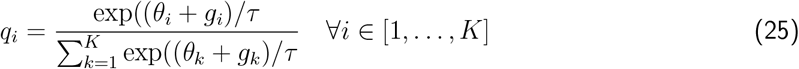

where *g*_*i*_ ∼ Gumbel(0, 1) is independent and identically distributed (i.i.d) noise samples drawn from the standard Gumbel distribution^20^. The state probabilities *α*_*t*_ are then given by applying softmax function *ξ* to the logits: 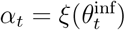.

**Secondly**, assuming that the data points are independent over time and only depend on the state active at the corresponding time point, we factorise the **likelihood** as

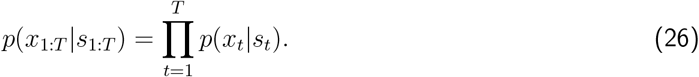

Each likelihood *p*(*x*_*t*_|*s*_*t*_) is modelled as a multivariate Gaussian distribution by

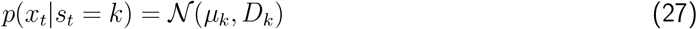

where *µ*_*k*_ and *D*_*k*_ are learnable parameters representing the mean vector and covariance matrix for each latent state *k*.

Now, substituting Equations (21) and (26) into Equation (20), our LL term becomes

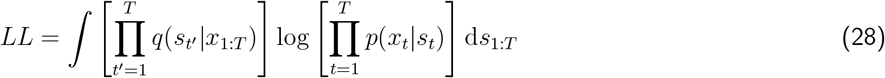

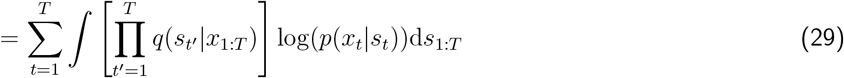

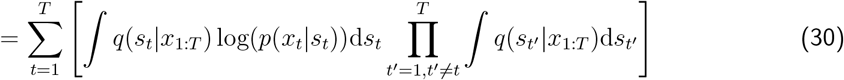

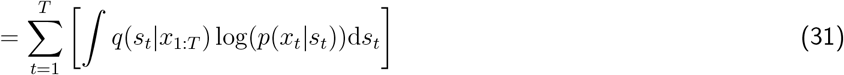

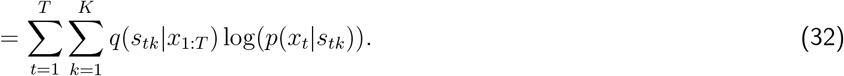

It is worthwhile to note that, for each time point, the LL term is a weighted sum (i.e., mixture) of the log probabilities under the state-specific multivariate normal distributions. In this way, the posterior from the **encoder** and the likelihood from the **decoder** are combined. During training, as the Gumbel-Softmax temperature *τ* is annealed towards 0 (see Section 2.1.2), the posterior state distribution will become increasingly categorical, and the mixture will become more like a hard assignment.

#### A.2.2 KL Divergence Loss

Considering the KL term

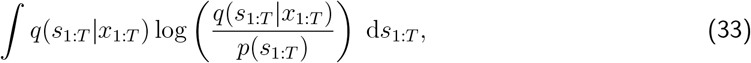

we first assume that the state at each time point is independent and only depend on the previous states. The **prior** is therefore factorised as

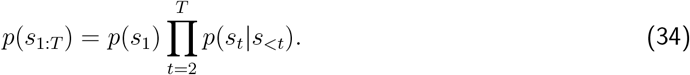

Substituting the factorised prior into Equation (33) and leveraging the mean field approximation in Equation (21), we now have

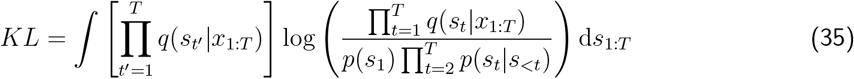

and split the logarithm to get

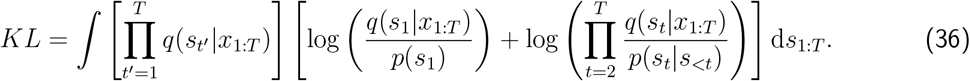

Because the first logarithm is positive and clipping it would keep the term proportional to the original value, we exclude it and approximate the KL divergence as

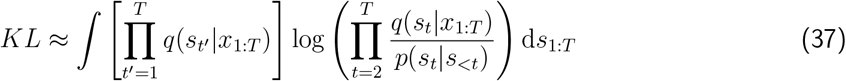

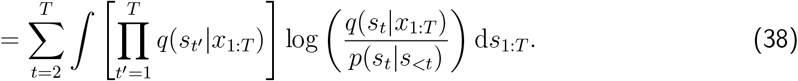

Next, for each term in the summation, we factorise the integral:

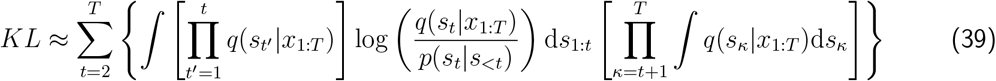

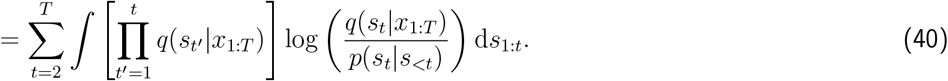

Furthermore, denoting the integral over d*s*_*t*_ by

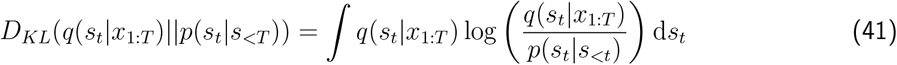

allows us to rewrite the KL term as

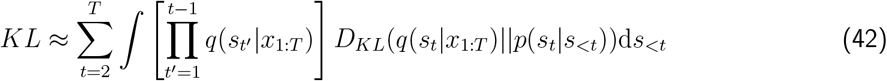

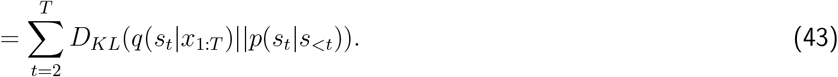

To calculate this, we can use the Monte Carlo estimation such that

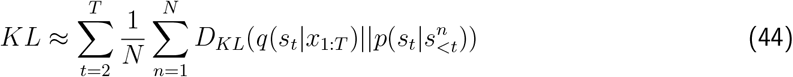

where *N* is the number of samples, and 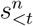 denotes the *n*^th^ sample from the posterior distribution *q*(*s*_*t*_|*x*_1:*T*_ ) at previous time points. For computational simplicity, *N* = 1 was used^21^. Therefore, KL is further approximated as

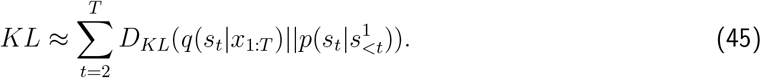

As explained in Section 2.1.2, the prior 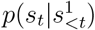 is modelled as a categorical distribution

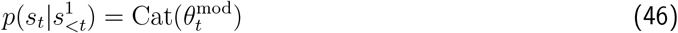

with 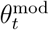 inferred using a unidirectional LSTM network (i.e., the model RNN) in the **regularization module** (see Equation (3)); the posterior *q*(*s*_*t*_|*x*_1:*T*_ ) is taken from the **encoder** as derived above.

#### A.2.3 Total Loss

Combining the LL and KL loss terms, our total loss ℒ is their sum

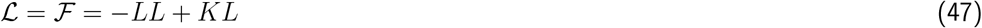

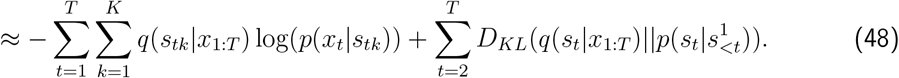

and we minimise this loss when training DyNeStE.

### A.3 Model Training Details and Key Hyperparameters

#### A.3.1 Additional Techniques for Model Training

To facilitate faster and more stable convergence of DyNeStE, we adopted several additional training strategies. These techniques were designed to mitigate common issues found when training a VAE with RNNs and discrete latent variables.

##### Adaptive Optimisation Strategies

Three adaptive mechanisms are incorporated into the training process:

- **KL annealing:** When training a VAE with RNNs, the KL divergence term can dominate the loss and force the model to drive the variational posterior distribution close to the prior, resulting in a negligible KL loss (Bowman et al., 2015). To address this problem, we apply KL annealing to the objective function:

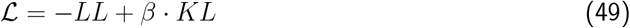

where *β* is a KL annealing factor that starts at 0 and gradually increases to 1 during training. This allows the encoder to first learn informative latent representations before aligning the variational distribution with the prior. The KL annealing schedule follows a hyperbolic tangent curve such that

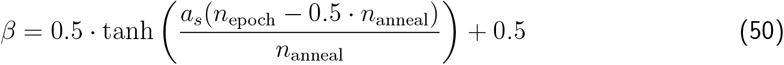

where *a*_*s*_ is the annealing sharpness, *n*_epoch_ is the total number of epochs, and *n*_anneal_ is the number of annealing epochs.
- **Gumbel-Softmax temperature annealing:** While Maddison et al. (2017) fixed the GumbelSoftmax temperature and reported minimal benefits for their experiments, Jang et al. (2017) employed a temperature schedule when training VAEs with the Gumbel-Softmax distribution. In our experiments, temperature annealing was essential for achieving stable and better model performance. We used an exponential decay schedule:

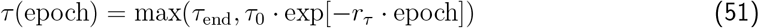

where *τ*_0_ and *τ*_end_ are the initial and final temperatures, respectively, and *r*_*τ*_ is the decay rate. This schedule allowed the model to learn a continuous posterior at the beginning and gradually transition into a more categorical distribution.
- **Learning rate scheduling:** We applied an exponential learning rate decay to accelerate training in the beginning and enable finer adjustments towards the end:

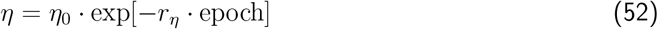

where *η*_0_ is the initial learning rate and *r*_*η*_ is the decay rate. In practice, we only begin decaying the learning rate after KL annealing is complete (i.e., *β* = 1). Note that this schedule was applied for HMM training as well.

##### Gradient Clipping

Although the aforesaid adaptive strategies improve convergence speed and stability, simultaneously varying multiple parameters (*β, τ*, and *η*) can make the optimisation landscape more complex. For a categorical model with discrete latent variables like DyNeStE, the loss surface tends to become increasingly noisy and non-smooth over time, especially as the Gumbel-Softmax temperature is annealed.

In certain model runs, we observed sporadic gradient explosions that destabilised training. To mitigate this, we constrained the gradient norm to a fixed threshold, preventing excessively large updates. While gradient clipping was unnecessary in our simulated experiments, it was crucial when training on real datasets.

##### Multi-start Initialisation

Since DyNeStE is trained using stochastic gradient descent, all trainable parameters are initialised randomly. The initialisation scheme for these parameters is as follows:

- **Weights and biases of the model and inference RNN:** Initialised using Glorot initialisation (Glorot & Bengio, 2010).
- **Layer normalisation parameters:** Layer normalisation (Ba et al., 2016) is applied between each RNN layer and subsequent dense layer in both the model and inference RNNs. Its learnable shift and scale parameters are initialised to zeros and ones, respectively.
- **Weights and biases of the dense layers:** Initialised using Glorot initialisation.
- **State means and covariances:** Mean vectors are fixed to zero, and covariance matrices are initialised using flattened Cholesky factors of identity matrices, with a Gaussian error added to the diagonal.

Because the objective function of DyNeStE is non-convex, its optimisation process is sensitive to parameter initialisation. We have observed that different random initialisations can result in high run-to-run variability, with a model converging to different local minima across runs. To mitigate this issue, we employed a multi-start initialisation strategy.

In this approach^22^, the model parameters are re-initialised multiple times. For each initialisation, the model is trained for a small number of epochs, after which the loss is evaluated. The initialisation that yields the lowest loss is selected for the main training. During this initialisation stage, neither the Gumbel-Softmax temperature nor the KL annealing factor is annealed. Empirically, we find that this choice yields improved model performance.

##### Observation Model Regularisation

When training DyNeStE on real MEG data, we apply an additional regularisation to the state covariance matrices of the observation model. Specifically, we impose an inverse Wishart prior on each state covariance *D*_*k*_, adding its negative log-density to the total loss:

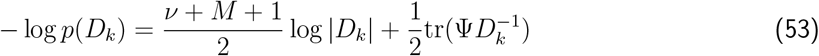

where *M* is the number of channels, *ν > M* − 1 is the degrees of freedom, | · | denotes the matrix determinant, and Ψ is a symmetric positive-definite scale matrix. The first term penalises large determinants, preventing covariances from inflating arbitrarily; the second term penalises small or ill-conditioned covariances, preventing them from shrinking toward singularity that could artificially increase the likelihood. This prior encourages each *D*_*k*_ to remain close to a target scale and shape specified by Ψ and *ν*, avoiding both excessive stretching and degeneration in extreme directions. The same covariance regularisation is applied for the HMM.

With the exception of the Gumbel-Softmax temperature annealing, all of these techniques have been applied in prior models; see DyNeMo (Gohil et al., 2022) and M-DyNeMo (Huang et al., 2025) for additional discussions on methodological details.

#### A.3.2 Model Training Curves

The loss curves of DyNestE and the HMM, trained on the Nottingham MEGUK dataset, are shown below. Each curve corresponds to the best-performing run selected from ten model runs.

**Figure A2:**
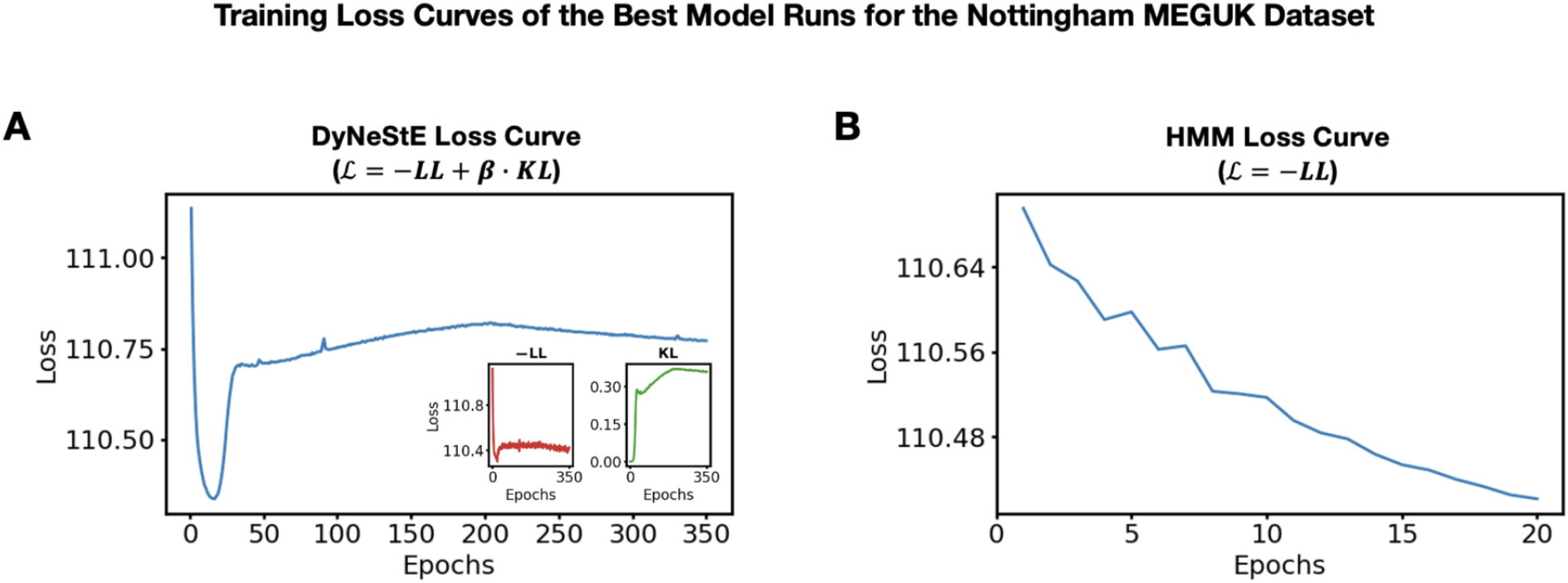
Training loss curves of DyNeStE and HMM. **(A)** Loss curve of DyNeStE (blue), where the loss is the variational free energy. The inset shows the negative log-likelihood (red) and KL divergence (green) terms separately. **(B)** Loss curve of the HMM (blue), where the loss is the negative log-likelihood.

To assess the stability of each model and characterise its variability across training runs, we extracted the final training losses from all ten runs and visualised them as a box plot. Note that the loss values of DyNeStE and the HMM are not directly comparable, as they are optimised using different objective functions.

**Figure A3:**
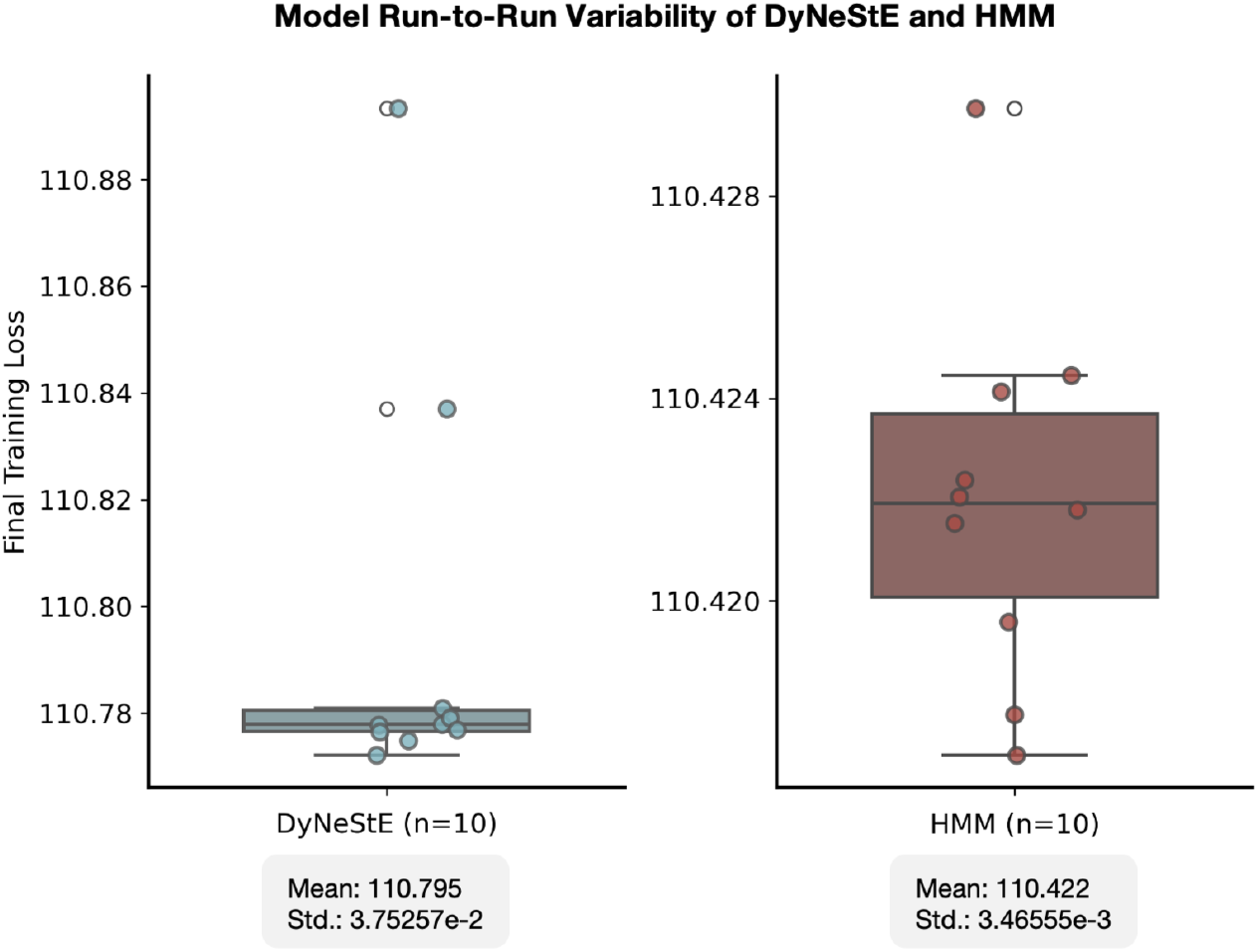
Run-to-run variability of final training loss for DyNeStE and HMM. Distributions of final training loss across 10 independent runs are shown as box plots for DyNeStE (left) and HMM (right). Coloured circles indicate loss values from individual runs, while open circles denote outliers. The mean and standard deviation of the final training losses for each model are reported under the corresponding box plot.

#### A.3.3 Model Hyperparameters

The hyperparameters of DyNeStE and the HMM used to train the models on the simulated and real MEG datasets are summarised in the tables below:

**Table 1:**
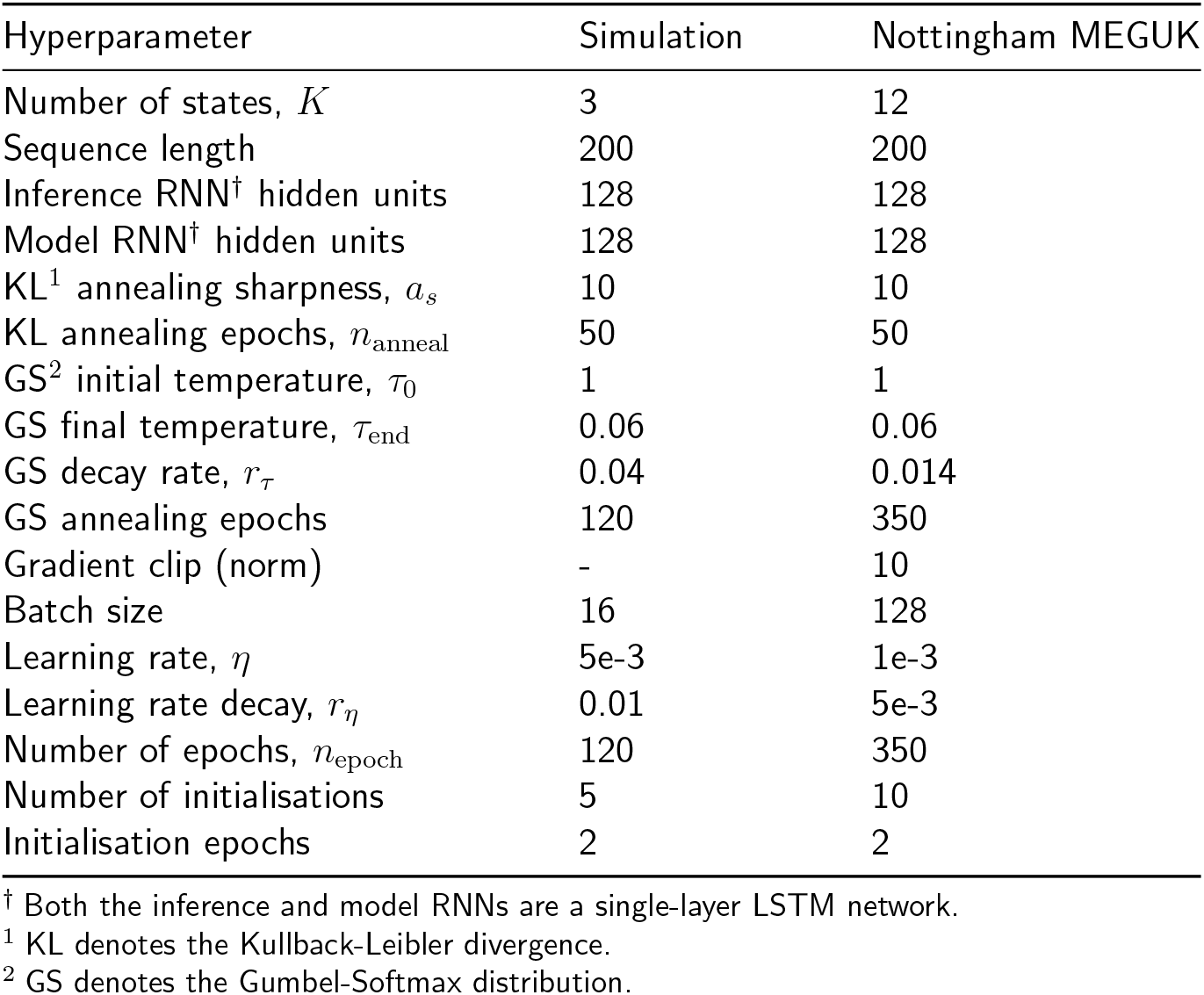
(DyNeStE) Hyperparameters for Simulation and Real MEG Data.

**Table 2:** (TDE-HMM) Hyperparameters for Simulation and Real MEG Data.

| Hyperparameter | Simulation | Nottingham MEGUK |
| --- | --- | --- |
| Number of states, $K$ | 3 | 12 |
| Sequence length | 200 | 200 |
| Batch size | 16 | 256 |
| Learning rate, $\eta$ | 0.01 | 0.01 |
| Learning rate decay, $r_\eta$ | 0.1 | 0.1 |
| Number of epochs, $n_{\text{epoch}}$ | 20 | 20 |
| Number of initialisations | 5 | 10 |
| Initialisation epochs | 2 | 2 |

Note that we set *K* = 12 for the real MEG datasets to allow direct comparison with previous studies that employed 12-state HMMs as their model specification (Higgins et al., 2021; van Es, Higgins, et al., 2025). For the split-half reproducibility analysis in Fig. A6, the number of states was systematically varied across *K* ∈ [4, 6, 8, 10].

In addition, the sequence length is chosen such that DyNeStE and HMM models have access to the same temporal context, ensuring a fair comparison. It has been tuned to optimise model performance under available GPU resources and compute time. For DyNeStE, the sequence length must be set to balance training stability and memory constraints against the need for a sufficiently long temporal window to capture long-range dependencies.

### A.4 Dataset Details

The subject demographic information for the Nottingham MEGUK dataset is summarised in the figure below:

**Figure A4:**
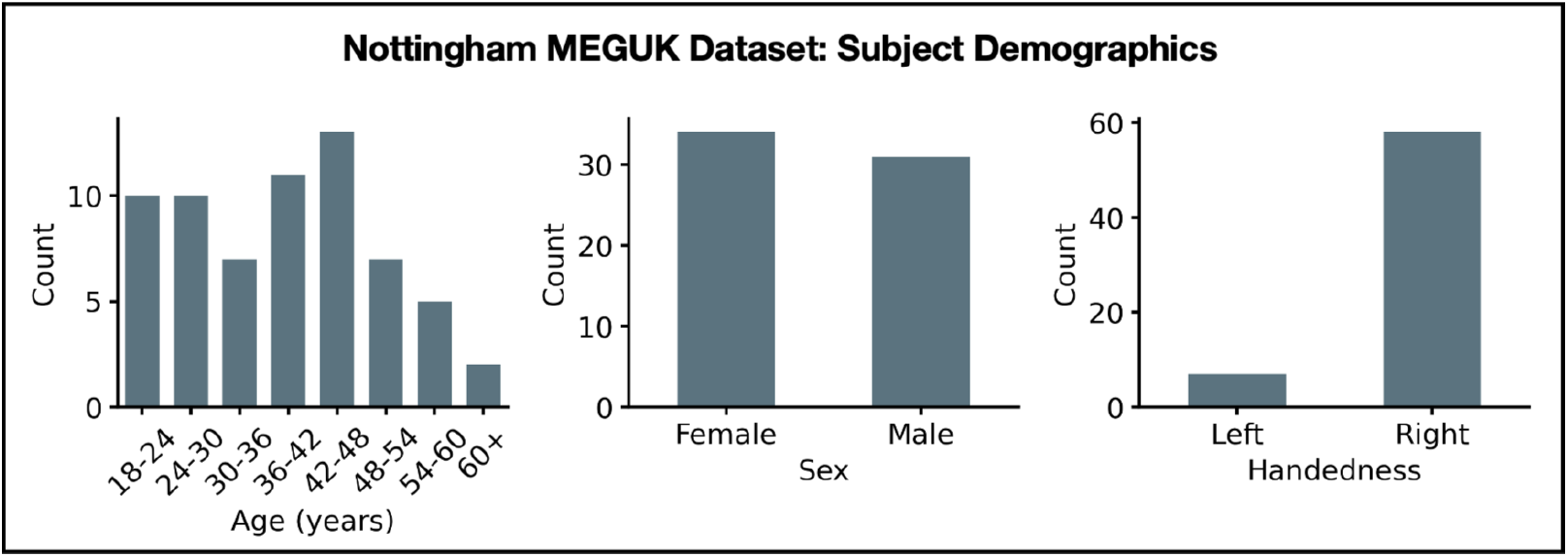
Nottingham MEGUK dataset subject demographics. Bar graphs depicting the histogram of participant ages (left), sexes (middle), and handedness (right).

### A.5 Additional Figures for the Nottingham MEGUK Dataset

Here, we append additional results on the Nottingham MEGUK dataset.

**Figure A5:**
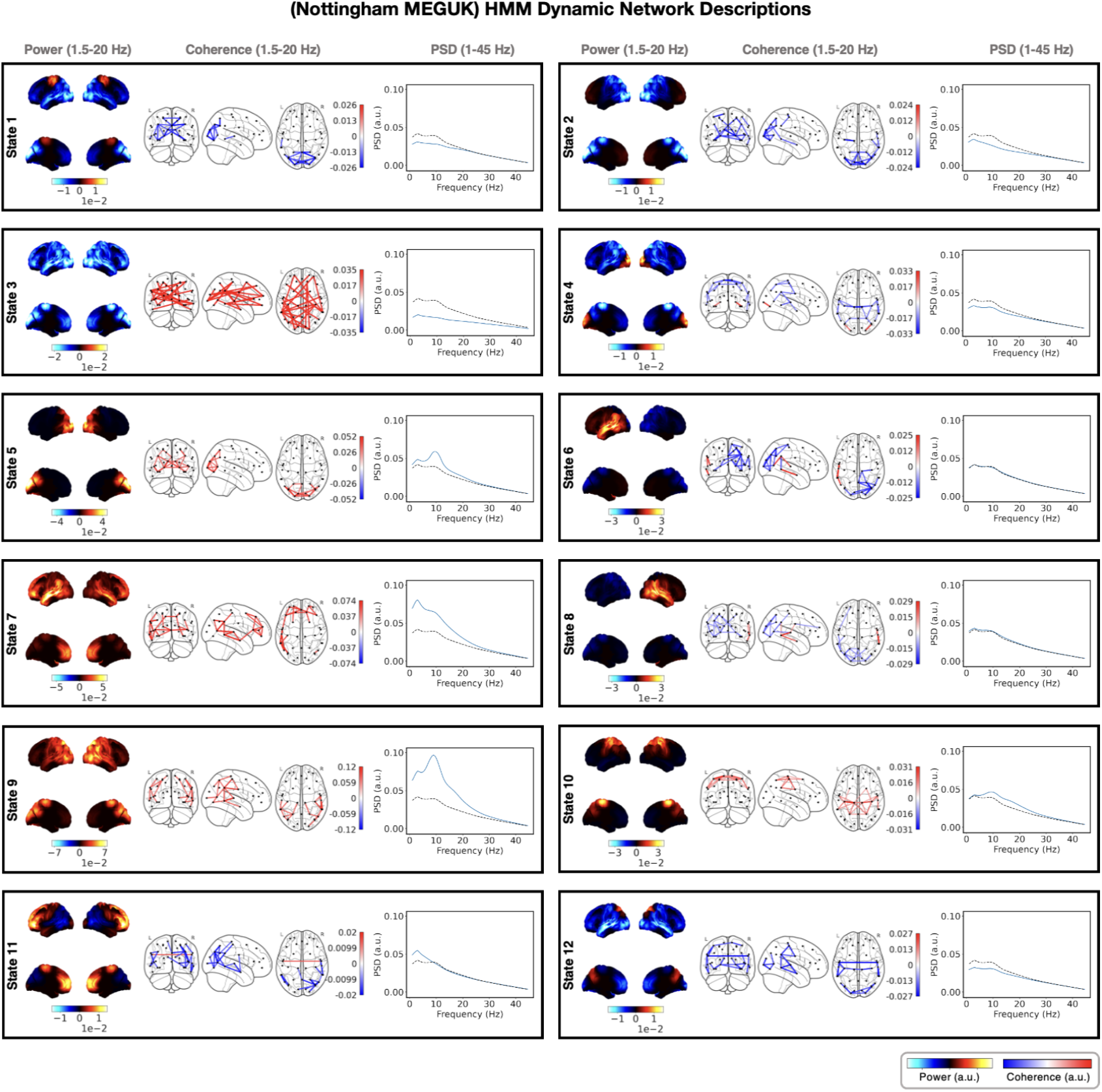
Dynamic resting-state MEG networks inferred by HMM. Using the HMM, twelve RSN states were identified from MEG recordings of 65 subjects. Each panel presents network descriptions of one state: the group-level power map (left), FC network (middle), and parcel-averaged PSD (right). The power maps display lateral and medial cortical surfaces at the top and bottom, respectively. The FC networks illustrate edges with the top 3% coherence values (regardless of sign). Both power maps and FC networks are shown relative to their average across all states. The PSD of each state (blue) is plotted alongside the state-averaged PSD (black dotted line).

**Figure A6:**
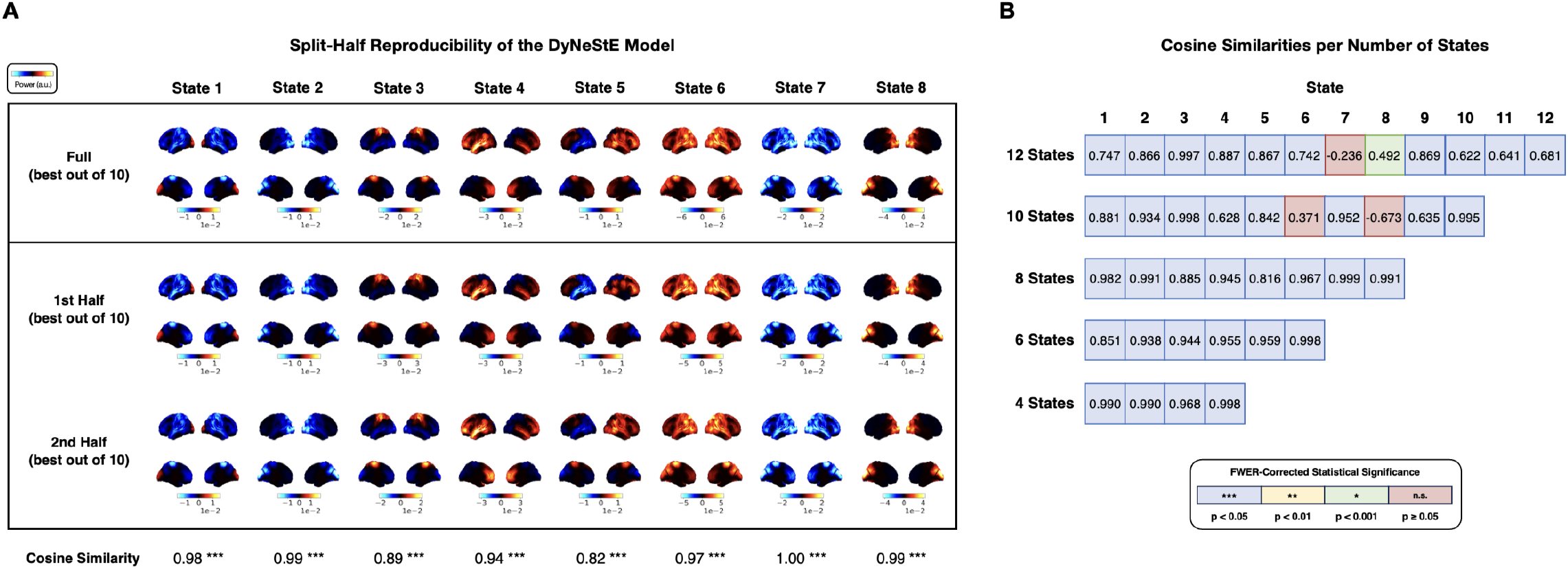
Split-half reproducibility of DyNeStE across different number of states. DyNeStE was trained on two split-halves of the Nottingham MEGUK dataset with varying numbers of states as its hyperparameter, selecting the best model (out of 10 runs) for each subset. **(A)** Group-level power maps of 8 states inferred from the full dataset (top), first split-half (middle), and second split-half (bottom), each displayed relative to the mean power across all states. Cosine similarities between split-half power maps are reported for each state. **(B)** Cosine similarities between split-half power maps are shown for different number of states. Because the model was fitted with varying state numbers, the resulting states correspond to different brain networks, and their ordering does not necessarily align across model fits. Statistical significance was assessed using a max-statistic permutation test over the states (***: *p <* 0.001, **: *p <* 0.01, *: *p <* 0.05, n.s.: non-significant). Note that the test controls for family wise-error rate (FWER) by constructing the null distribution of max statistic over all states.

**Figure A7:**
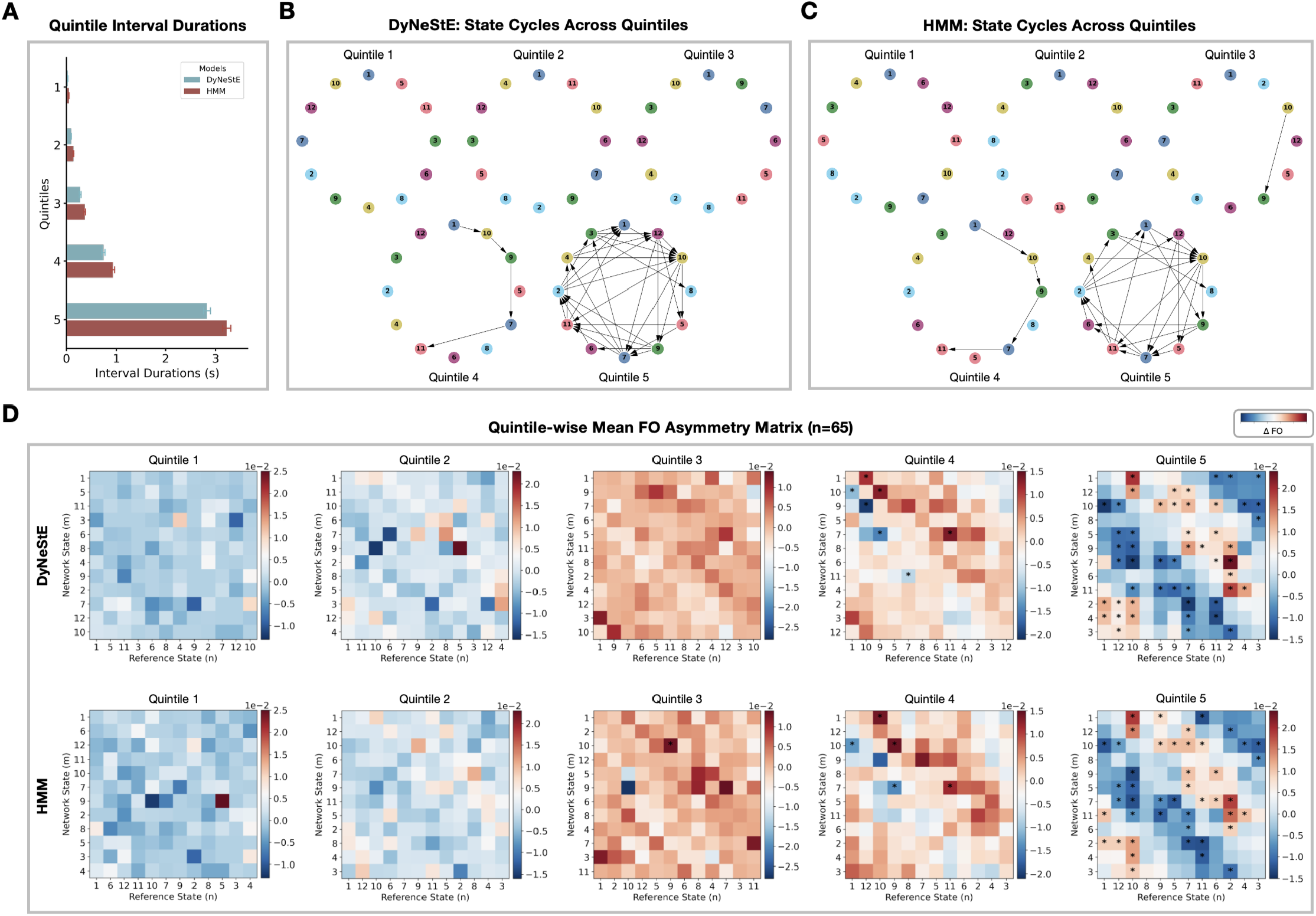
Quintile-wise cyclical organisation of state activations in observed data. **(A)** Distributions of inter-state interval durations, extracted from the inferred state time courses, are shown across quintiles. Mean values and standard errors are depicted as bar plots and error bars, respectively, for DyNeStE (blue) and the HMM (red). **(B)** Quintile-specific state cycles derived from the DyNeStE-inferred state time courses. **(C)** Quintile-specific state cycles derived from the HMM-inferred state time courses. **(D)** Group-averaged FO asymmetry matrices for each quintile, shown for DyNeStE (top) and the HMM (bottom). Asterisks mark statistically significant edges identified using a paired-samples t-test (*p <* 0.05, Bonferroni-corrected with *n* = 132 state pairs).

### A.6 Additional Figures for the Replay Dataset

Lastly, we append additional results on the Replay dataset.

**Figure A8:**
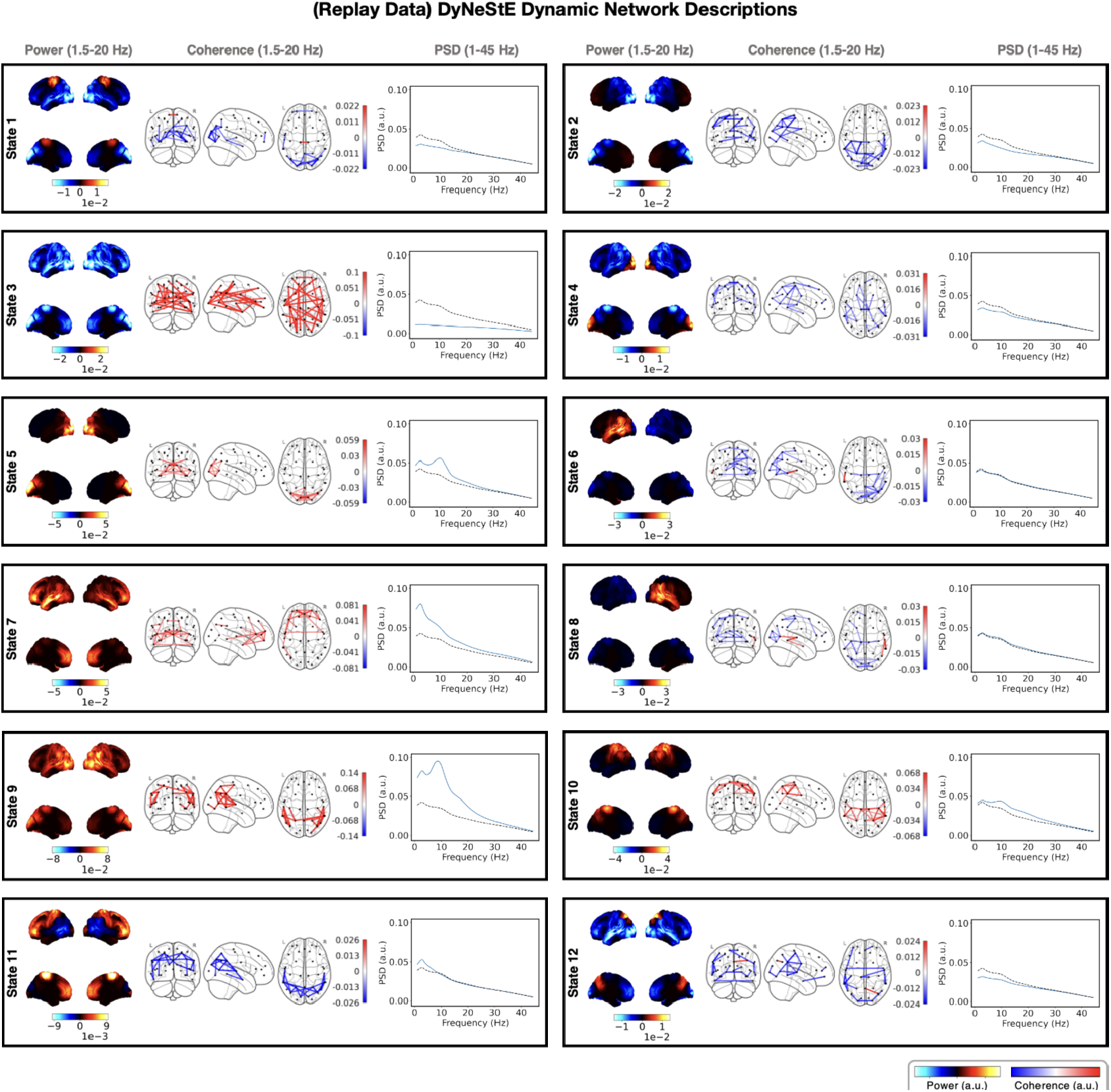
Dynamic resting-state MEG networks inferred by DyNeStE. Twelve RSN states were identified from MEG recordings comprising 42 sessions (21 subjects, two sessions each) in the Replay dataset using DyNeStE. Each panel presents network descriptions of one state: the group-level power map (left), FC network (middle), and parcel-averaged PSD (right). The power maps display lateral and medial cortical surfaces at the top and bottom, respectively. The FC networks illustrate edges with the top 3% coherence values (regardless of sign). Both power maps and FC networks are shown relative to their average across all states. The PSD of each state (blue) is plotted alongside the state-averaged PSD (black dotted line).

**Figure A9:**
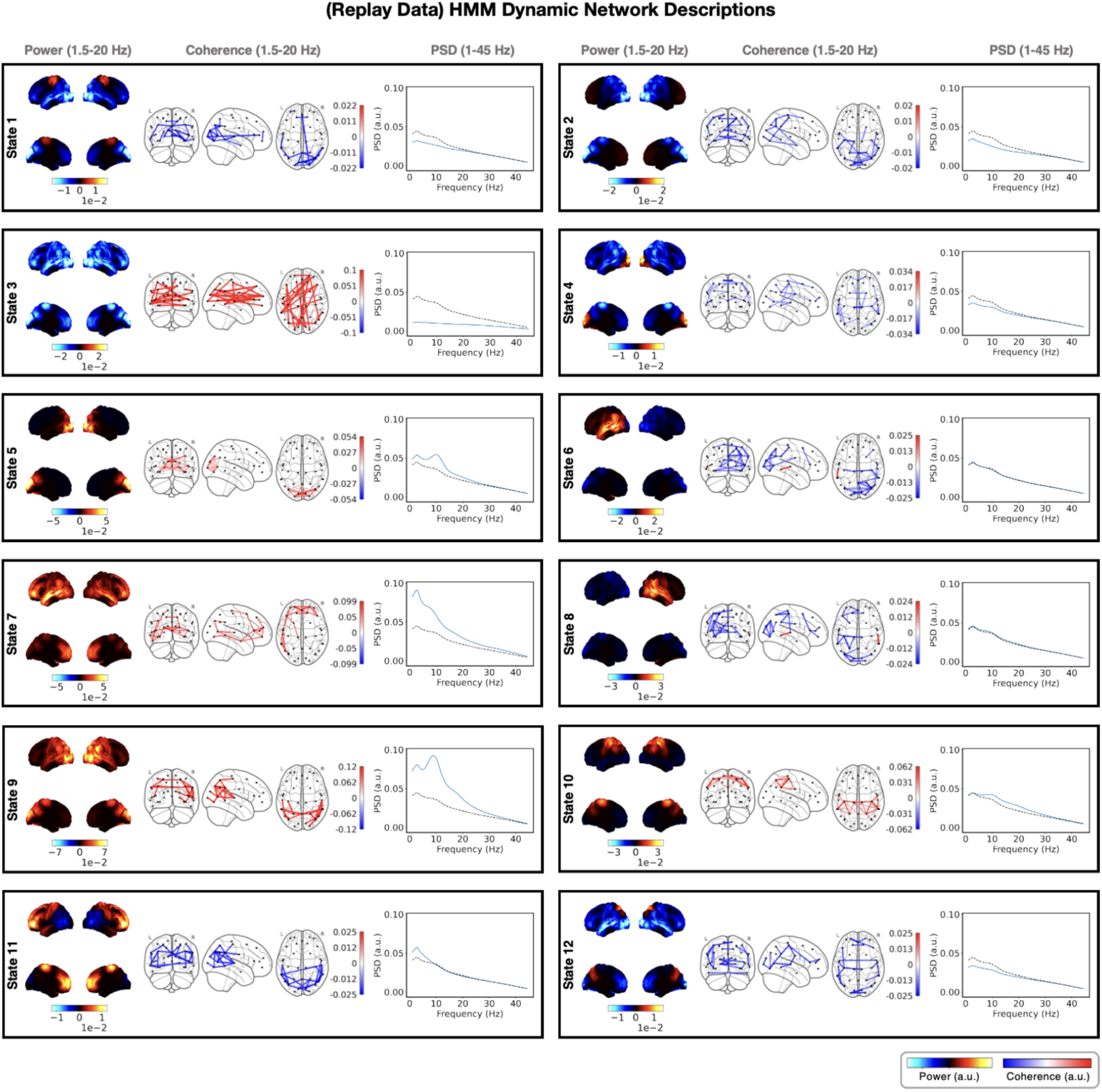
Dynamic resting-state MEG networks inferred by HMM. Twelve RSN states were identified from MEG recordings comprising 42 sessions (21 subjects, two sessions each) in the Replay dataset using the HMM. Visualisations are presented in the same format as in Figure A8.

## Footnotes

1 Formally, this property is described as the HMM following a first-order Markovian process.

2 This approach has also been adopted in previous models, including DyNeMo (Gohil et al., 2022), Multi-dynamic DyNeMo (M-DyNeMo) (Huang et al., 2025), Time-Delay Embedded HMM (TDE-HMM) (Vidaurre et al., 2018), and HMM with Integrated Variability Estimation (HIVE) (Huang et al., 2024).

3 For conciseness, we hereinafter denote *s*_1:*t*−1_ as *s*_*<t*_.

4 We refer to the inferred networks as *modes* rather than *states* to emphasise their continuous nature. Unlike states, multiple network modes can be active simultaneously at a given time point.

5 It should be noted that while the states *s*_*t*_ are learned as probability distributions, the logits that parametrise this distribution, as well as other model parameters, are learned as point estimates.

6 If the state-specific mean vector *µ*_*k*_ were to be learnt instead of being set as a zero vector, it can take on any value and can therefore be treated as free parameters.

7 In practice, feeding the entire time series to an LSTM is infeasible due to GPU memory constraints and the vanishing gradient problem. Therefore, the data is divided into shorter sequences, and at each optimisation step during training, a mini-batch of shuffled sequences is used to compute the loss and gradients. See Section 2.5 and Equation 9 in Huang et al. (2025) for more details.

8 Here, applying PCA is theoretically preferable than simply using descriptive statistics such as the maximum or mean. While such measures are computationally efficient, they capture only limited aspects of the data. In contrast, PCA identifies components that explain maximal variance, taking variability of the data distribution into account.

9 The use of subject-specific sMRI is recommended when available. A previous study has shown that employing subject-specific sMRI, rather than a template, for source reconstruction can improve age-group classification accuracy for both EEG and MEG data, when resting-state network features extracted using HMMs are used as input (Cho et al., 2024). This implies that models for brain-state analysis can capture and may benefit from subject-specific anatomical information retained during source reconstruction.

10 Note that the different choices in data preparation, including the TDE window length and the number of principal components, result in data with different oscillatory properties. Care must be taken to use sufficiently many temporal lags and components to preserve both low- and high-frequency activity.

11 For brevity, we refer to the TDE-HMM model as HMM throughout the remainder of this paper.

12 This corresponds to 102.4 seconds at a sampling rate of 250 Hz. In the subsequent post-hoc analyses, the simulated data was trimmed to match the generated data length.

13 Nonetheless, it is important to note that different FC measures emphasise distinct aspects of brain connectivity and may lead to observable differences in the resulting network patterns. For instance, orthogonalised amplitude envelope correlation (AEC) has been used in our prior work. However, it primarily captures co-modulation of oscillatory amplitudes and is thus better suited to static FC analyses, where network-level coordination at slower timescales is stronger (cf. (Colclough et al., 2016)). Accordingly, the choice of FC measure should depend on the data, model, and hypothesis of one’s interest.

14 The state time courses correspond to the maximum a posteriori estimate obtained from the inferred posterior distribution (see Section 2.3).

15 In prior work, it was shown that when HMM state time courses are simulated from the transition probability matrix for durations 100 times longer than the observed data, both FO asymmetry and state cycles start to emerge from the generated state time courses (see Figure S4 in van Es, Higgins, et al. (2025)).

16 The absence of sMRI scans can lead to less accurate source-reconstruction and, consequently, imprecise spatial estimation (see Section 2.3).

17 It is important, however, to distinguish inference from the generative capacity of a model. As emphasised in the *Results*, the ability of the HMM to identify long-term dynamics during *inference* does not mean it has learnt long-range temporal dependencies through its generative model parameters (i.e., via its transition probability matrix). This is a known limitation of the HMM. As illustrated in Figures 6-9, the inferred HMM failed to *generate* data with long-term dynamics, demonstrating its limited representational capacity.

18 Nonetheless, caveats do exist. Iterative training on recursively generated data can cause model degradation (Shumailov et al., 2024), and partially synthetic datasets may still retain links to original participant records, limiting their ability to guarantee privacy (Zhang et al., 2022).

19 An epoch refers to a single iteration over the entire dataset.

20 Each *g*_*i*_ is drawn using the Gumbel-Max trick (Gumbel, 1954; Maddison et al., 2014), having *g*_*i*_ = − log(− log(*U*_*i*_)) with *U*_*i*_ ∼ Uniform(0, 1).

21 Empirically, taking *N >* 1 did not enhance the model performance noticeably and does not seem to increase the accuracy of estimation significantly.

22 For the HMM, we adopt a slightly modified initialisation procedure. State time courses are randomly sampled, and the state-specific mean vectors and covariance matrices are computed from these samples. These empirically estimated means and covariances are then used to initialise the corresponding model parameters for each state. All remaining parameters are initialised following the standard scheme described above.

## Notes

### Competing Interest Statement

The authors have declared no competing interest.

### Summary of Updates

Improved the simulation analyses by incorporating real data-driven covariance matrices; Figure 2 revised; Figure A3 added; expanded and elaborated methodological choices

